# Genome-wide analysis of *Schistosoma mansoni* reveals population structure and praziquantel drug selection pressure within Ugandan hot-spot communities

**DOI:** 10.1101/2022.01.25.477652

**Authors:** Tushabe John Vianney, Duncan J. Berger, Stephen R. Doyle, Geetha Sankaranarayanan, Joel Serubanja, Prossy Kabuubi Nakawungu, Fred Besigye, Richard E. Sanya, Nancy Holroyd, Fiona Allan, Emily L. Webb, Alison M. Elliott, Matt Berriman, James A. Cotton

**Author notes:** **Author Contributions** TJV: Formal analysis, Investigation, Writing – Original Draft Preparation, Writing – Review & Editing DJB: Formal analysis, Investigation, Writing – Review & Editing SRD: Formal analysis, Investigation, Writing – Review & Editing GS: Investigation, Resources, Writing – Review & Editing JS: Investigation, Resources, Writing – Review & Editing KP: Investigation, Writing – Review & Editing FB: Investigation, Writing – Review & Editing RES: Conceptualisation, Investigation, Resources, Writing – Review & Editing NH: Project Administration, Writing – Review & Editing FA: Investigation, Methodology, Resources, Writing – Review & Editing ELW: Conceptualization, Formal analysis, Methodology, Writing – Review & Editing AE: Conceptualization, Project Administration, Supervision, Funding Acquisition, Writing –Review & Editing MB: Conceptualization, Project Administration, Supervision, Funding Acquisition, Writing – Review & Editing JAC: Formal analysis, Supervision, Writing – Original Draft Preparation, Writing – Review & Editing.

## Abstract

Populations within schistosomiasis control areas, especially those in Africa, are recommended to receive regular mass drug administration (MDA) with praziquantel (PZQ) as the main strategy for controlling the disease. The impact of PZQ treatment on schistosome genetics remains poorly understood, and is limited by a lack of high-resolution genetic data on the population structure of parasites within these control areas. We generated whole-genome sequence data from 174 individual miracidia collected from both children and adults from fishing communities on islands in Lake Victoria in Uganda that had received either annual or quarterly MDA with PZQ over four years, including samples collected immediately before and four weeks after treatment. Genome variation within and between samples was characterised and we investigated genomic signatures of natural selection acting on these populations that could be due to PZQ treatment. The parasite population on these islands was more diverse than found in nearby villages on the lake shore. We saw little or no genetic differentiation between villages, or between the groups of villages with different treatment intensity, but slightly higher genetic diversity within the pre-treatment compared to post-treatment parasite populations. We identified classes of genes significantly enriched within regions of the genome with evidence of recent positive selection among post-treatment and intensively treated parasite populations. The differential selection observed in post-treatment and pre-treatment parasite populations could be linked to any reduced susceptibility of parasites to praziquantel treatment.

**Author summary:** Schistosomiasis is caused by parasitic helminths of the genus *Schistosoma*. *Schistosoma mansoni* is the primary cause of intestinal schistosomiasis, a devastating and widespread parasitic infection that causes morbidity, death and socio-economic impact on endemic communities across the world and especially sub-Saharan Africa. Using whole-genome sequencing, we were able to elucidate the parasite population within Lake Victoria island fishing communities in Uganda which are among the major hotspots for schistosomiasis. We further assessed genetic markers that might be linked to recent observations concerning reduced susceptibility to praziquantel, the major drug used in the treatment of this disease. Whole-genome data on the population genetics of *S. mansoni* in an African setting will provide a strong basis for future functional genomics or transcriptomic studies that will be key to identifying drug targets, improving existing drugs or developing new therapeutic interventions.

## Introduction

Schistosomiasis – also known as Bilharzia after its discoverer Theodor Bilharz [1] – is a neglected tropical disease that affects about 250 million people worldwide, most of whom live in low and middle-income countries (LMICs) [2]. To treat schistosomiasis, praziquantel (PZQ) is used for preventative chemotherapy by mass drug administration (MDA)[3] and has been used globally to treat schistosome infections since 1979 [4]. In Uganda, the ongoing use of PZQ in MDA started between 2002 and 2003 [3, 5]. The objective of MDA in these settings has historically been to reduce the prevalence and intensity of infection and hence pathology; cure and elimination are not expected in the absence of additional interventions such as improving sanitation and snail control [6, 7]. In the World Health Organisation 2021-2030 the goal has been set of reducing the proportion of people with high-intensity infections to < 1% and thereby to eliminate schistosomiasis as a public health problem in all countries in sub-Saharan Africa by 2030 [8]. The expectation is that this will be achieved primarily by increasing the frequency and coverage of treatment with PZQ – the sole drug commonly used for schistosomiasis MDA –which could inadvertently increase drug selection pressure on parasite populations.

There is a growing body of evidence that MDA programmes may affect how parasite populations respond to treatment, for example, through reduced efficacy of PZQ in lowering egg output in communities that have received multiple rounds of PZQ MDA [9, 10], but there is little evidence that this is a widespread phenomenon [11, 12]. Reduced genetic diversity of parasite populations has also been associated with reduced susceptibility of the parasites to PZQ [13], with reports from Senegal having earlier linked such outcomes to potential drug resistance [14]. The development of drug resistance in natural populations would be a major health concern. Furthermore, *in vitro* studies have shown that resistance to PZQ can be selected for in *S. mansoni* [13, 15–17]. There is growing interest in understanding the impact of continued PZQ monotherapy on the parasite genome in order to detect the potential development of resistance to this drug as early as possible [18, 19], and understand the mechanism(s) of resistance. One clue to resistance can come through understanding the mode of action of a drug. The activity of PZQ has not been clearly understood, but recent findings suggest that the drug activates schistosome Ca^2+^-permeable transient receptor potential (TRP) channel (Sm.TRPMPZQ) [20], hence making it the primary target for PZQ action on schistosomes. Recently, a genetic cross involving a schistosome line experimentally selected for PZQ resistance identified this TRP channel as likely responsible [21], but it is not yet clear that this locus is involved in variation in PZQ efficacy in the field. Other candidate genes have been proposed, for example the *S. mansoni* P-glycoprotein (smdr2), which shows increased protein expression in male worms following exposure to sub-lethal doses of PZQ [16]. Susceptibility of the parasites to PZQ might involve multiple interactions between the drug, the parasite, and the respective host.

Collecting high-resolution genetic data from parasite populations under drug selection pressure may lead to new insights into the mode of action of PZQ or the mechanism of potential resistance to the drug. Furthermore, population genetic data from parasite populations will also give insights into the population biology of the parasite. This is vital for understanding schistosomiasis epidemiology, transmission, disease severity and why certain communities might respond better to treatment than others, especially within regions where drug selection pressure is being applied[22]. While lower-resolution markers have been extensively used (e.g.[23, 24]), much of our detailed understanding of schistosome population genetics has come from studies using microsatellite markers to describe the genetic structure of populations of *S. mansoni* [25–27] and other schistosome species [28, 29]. This work has revealed genetic differentiation between parasite populations that are geographically separated (e.g [30–33]), but panmictic populations and very high within-host diversity within disease foci (e.g. [34, 35]). The population genetics of African schistosomes has recently been reviewed [36]. Microsatellite markers have also been employed to investigate both basic questions about parasite biology (e.g. [37]) as well as more applied, operational questions about schistosome control [22]. In particular, a few studies have shown changes in genetic diversity of schistosomes with praziquantel MDA [4, 38, 39], but other studies have failed to find this effect [40] particularly with longer-term follow-up [41] suggesting any genetic response to treatment may be only temporary [42].

With their high levels of polymorphism, microsatellite loci are powerful molecular markers, but inevitably represent only a small proportion of the parasite genome. There is an increasing amount of genome-scale data available for schistosome populations. A number of studies have used exome capture [43] to describe introgression between *Schistosoma* species [44] and to study the historical demography of schistosomiasis in the Americas [45]. Restriction site-associated sequencing (ddRAD-seq) has been used to demonstrate strong genetic structure in remaining endemic hot-spots of *S. japonicum* transmission [46, 47]. While providing high-resolution data in a cost-effective way, these reduced-representation sequencing approaches have some drawbacks, for example in identifying small haplotype blocks from ancient introgression [44]. Whole-genome data gives a more comprehensive picture of genetic variation, including non-coding variation, and so has the potential to provide more insights into understanding the population genetics of this species. While reference genome assemblies are available for a number of schistosome species [48–54], large-scale genome-wide variation data is only available from one *S. mansoni* population [55], with a number of other populations and other species most being represented by relatively few specimens [53, 56–58]. Efforts in elucidating the parasite population genetic structure have proven very helpful in understanding drug resistance or transmission mechanisms in other parasite species: most notably in the malaria parasite *Plasmodium falciparum* [59, 60] for which very extensive genome data is available [61].

Within Uganda, the Lake Victoria Island Intervention Study on Worms and Allergy-related diseases (LaVIISWA) was a cluster-randomised clinical trial [62] examining the impact of intensive (quarterly) versus standard (annual) PZQ treatment. While the study was primarily designed to assess the impact of anthelmintic treatments on allergy-related outcomes, prevalence and intensity of *S. mansoni* was a secondary outcome with results suggesting a plateauing of infection, after an initial decline in intensively treated villages [63]. To assess this outcome, a pilot study in the fourth year of the LaVIISWA trial investigated cure rate and Egg Reduction Rate (ERR) [10]. A lower cure rate and ERR was seen among people receiving quarterly (intensive) treatment (n=61; cure rate 50.8%, 95% confidence interval (CI): 37.7% to 63.9%; ERR 80.6%, 95% CI: 43.8% to 93.7%) than in those receiving a single annual standard dose (n=49, cure rate 65.3%, 95% CI: 50.4% to 78.3%; ERR 93.7%, 95% CI: 84.9% to 97.7%) [10]. The WHO recommends an ERR of 90% for effective PZQ treatment [9, 64]. While the sample size available precluded finding compelling statistical evidence, these results are suggestive of the first signs of reduced efficacy of PZQ treatment in the more intensively treated population, and that the plateau in reduction of infection during the intervention study could be due to PZQ resistance. These islands thus represent a ‘hot spot’ in which high baseline prevalence [62] of schistosomiasis has persisted despite multiple years of treatment [10, 65].

Here, we sought to establish genome-wide data on the population genetics of parasites present in this study population, with the ultimate goal of assessing the effects of MDA on parasite genome evolution. We take advantage of the opportunity to investigate these in the context of a randomised intervention trial within a defined geographical area, allowing us to compare the effects of geographical isolation and treatment intensity on genetic variation in this population. By comparing samples taken immediately before and after a treatment round at the end of the LaVIISWA study in the two treatment arms and for multiple villages (the level of randomisation in the study), we can investigate whether the genetic impact of a single treatment dose varies with history of drug exposure. Building on the evidence that there may be differences in treatment efficacy between treatment arms, we investigated whether the signatures of natural selection across the genome differ with previous drug exposure. We also compare these data with recently published genomic data from other Ugandan *S. mansoni* populations.

## Methods

### Ethical considerations

This work was not expected to result in any harm to participants. Ethical approval was given by the Uganda Virus Research Institute (reference number GC127), the Uganda National Council for Science and Technology (reference number HS 1183) and the London School of Hygiene & Tropical Medicine (reference number 6187). As previously detailed [62], written informed consent was received from all adults and emancipated minors and from parents or guardians for children; additional assent was obtained from children aged ≥8 years.

### Sample selection and study site

Participants were selected from four villages each from the standard and intensive treatment arms from among the 27 study villages of Lake Victoria Island Intervention Study on Worms and Allergy-related diseases (LaVIISWA) trial [62, 63] at the end of its fourth year. The participants involved children and adults as previously described [62]. The villages in the standard arm received PZQ once a year while those in the intensive arm received PZQ four times a year during the LaVIISWA trial period. The standard villages sampled were Kakeeka, Kachanga, Zingoola and Lugumba. The intensive villages were Busi, Kitosi, Kisu and Katooke (Fig. 1).

**Fig 1.**
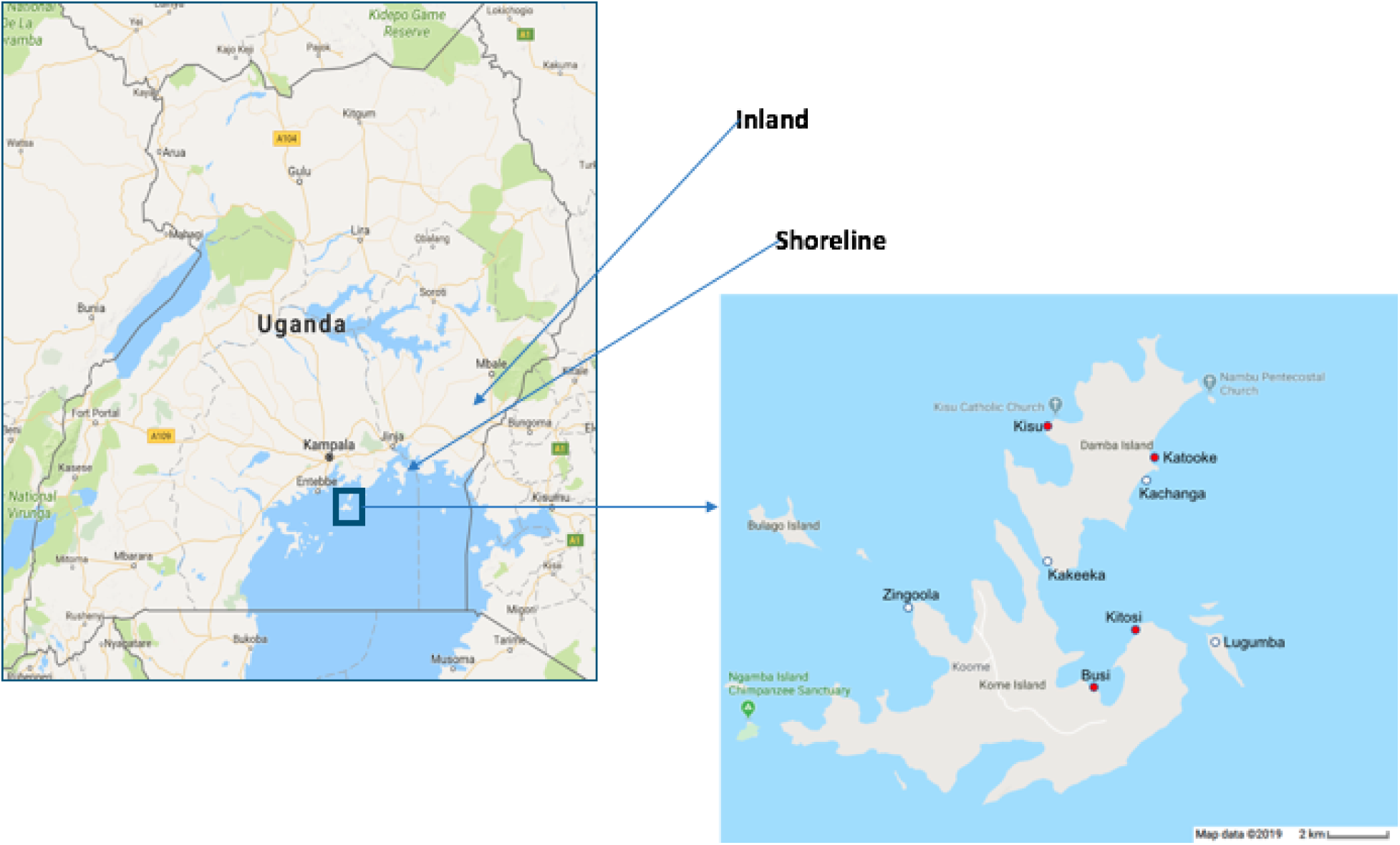
Location of sample sites within Uganda. Villages with white dots received standard (annual) intervention, those with red dots received intensive (quarterly) intervention. Outgroup samples were obtained from locations marked as inland and shoreline. Map data copyright 2019 Google.

Sample selection and collection was carried out as previously described in the parasitological survey [10]. The stool samples (collected from participants who tested positive for urine CCA) were processed for two Kato Katz slides as previously described [10] and miracidia hatching provided suitable material for DNA extraction. Participants were then treated, under observation, with a single dose of PZQ at 40 mg/kg (estimated by height pole), in accordance with the trial MDA procedures. Individuals whose pre-treatment sample tested positive for schistosome eggs by Kato Katz were followed up after four weeks and both Kato Katz and miracidia hatching were repeated. Miracidia hatching was carried out from each of these participants and the resultant miracidia were stored on Whatman FTA cards until DNA was extracted.

### Miracidia hatching

Miracidia hatching was carried out following previously described protocols [31]. In brief, the stool sample was homogenised through a metal sieve, then further washed and filtered using a Pitchford funnel assembly [66] consisting of a 40 μm sieve placed inside a 200 μm outer sieve. Stool samples were washed using deionised water (Rwenzori Bottling Company, Uganda). The concentrated *S. mansoni* eggs were transferred to a Petri dish in clean water and exposed to indirect sunlight to induce the hatching of miracidia. Hatching was performed in natural light (environmental conditions) with intervals of exposure to sunlight and cover depending on weather conditions. The time taken for miracidia to emerge varied between samples, so the Petri dishes were intermittently checked for the presence of miracidia for a maximum of 48 hours. Miracidia were picked in 1.5 – 5μl of water and then transferred to a second dish of deionised water to dilute bacterial contamination before being placed on Whatman indicating FTA cards (Qiagen) and left to dry. The FTA cards were wrapped in aluminium foil to keep them away from continued direct light and placed in ziplock bags with silica gel in a cardboard drawer.

### Whatman FTA DNA Extraction

DNA was extracted using a modified CGP buffer protocol as previously described [67, 68]. The individual spots containing miracidia were punched from the FTA cards using a 2 mm Harris micro-punch and placed in 96-well plates. Protease buffer was prepared using Tris-HCl pH 8.0 (30 mM), Tween 20 (0.5%), IGEPAL CA-630 (0.5%), protease (1.25 μg/ml; Qiagen cat #19155) and water. Digestion was done by adding 32 μl of the protease buffer to each of the wells on the 96- well plate containing the punched spots from the FTA cards. The plate was vortexed to mix and spun down before incubation at 50°C for 60 min, 75°C for 30 min. Miracidia lysates containing DNA were transferred to a new labelled plate and stored at 4°C until used.

### Library preparation and sequencing

DNA sequencing libraries were prepared using a protocol designed for library preparation of Laser Capture Microdissected Biopsy (LCMB) samples using the Ultra II FS enzyme (New England Biolabs) for DNA fragmentation as previously described [68]. The LCMB library preparation method is optimised for uniform, low-input samples. A total of 12 cycles of PCR were used to amplify libraries and to add a unique 8-base index sequence for sample multiplexing. The LCMB library preparation protocol is optimised for uniform, low input samples. A total of 174 samples were sequenced on two NovaSeq lanes, 108 on one lane and 66 on another lane. These 174 samples were chosen as having more than 10% of reads mapping to *S. mansoni* based on preliminary low-coverage genome sequencing of all 214 samples collected in the field.

### Mapping and SNP calling

The reads were mapped to the *S. mansoni* reference genome v7 (GCA_000237925.3) [54] using the BWA-MEM algorithm in Burrows-Wheel Aligner software (BWA) (VN:0.7.15-r1140) to produce SAM files which were then converted to BAM format using Samtools v1.14 . This version of the reference genome was modified to remove haplotypes in order to improve mapping accuracy, as previously described [55]. PCR duplicate reads were identified using Picard v1.92 [69] and flagged as duplicates in the BAM file.

SNP variants were called using the GATK Haplotype Caller (v4.1.4.1) to find sites that differ from the *S. mansoni* reference genome followed by variant QC to remove low confidence SNPs and regions of consistently poor calls. The SNPs were hard-filtered in GATK to remove SNP calls with the following parameters: QD) < 2.0; MQ < 40; FS) > 60.0; SOR > 3.0; MQRankSum < -12.5; ReadPosRankSum < -8.0. The variants were further filtered using vcftools_0.1.15 [70] to remove sites with high missingness (--max-missing 0.95), low minor allele frequency (--maf 0.01) and to retain only biallelic SNPs (--min-alleles 2 --max-alleles 2).

### Identification of population structure

The three islands on which the population structure was assessed were Koome, Damba and Lugumba in the Mukono district of Uganda. An outgroup made up of inland and shoreline samples was also included, consisting of 27 samples collected in a previous study [9] and for which whole-genome sequence data were recently published [55] from Tororo and Mayuge districts in Eastern Uganda. Tororo and Mayuge are approximately 120 km apart. Mayuge district is a shoreline district located about 100 km from Mukono district. Both districts are located in south eastern Uganda, with Tororo being the inland district (Fig. 1).

### Test for genetic differentiation

The fixation index (F_ST_) statistic was calculated between each of the villages across the different islands and treatment groups (standard, intensive, pre-treatment and post-treatment) to measure population differentiation due to genetic structure. The F_ST_ was calculated using vcftools (version 0.1.15) [55] on the vcf file containing biallelic filtered SNPs. Mean F_ST_ was calculated from genome-wide weighted F_ST_ values with 99% symmetric bootstrap confidence intervals calculated using R version 3.5.1 (2018-07-02). We fitted a gravity model as

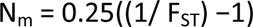

where N_m_ is an estimated number of migrants per generation, calculated from the F_ST_ between villages as:

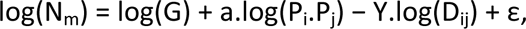

And G is the linear distance between the villages. In (P_i_.P_j_), P_i_ and P_j_ represent the population sizes of the two villages compared. Models were fitted using R version 4.02, with the MuMIn package v1.43 to assess model importance. Code for these analyses is available at https://github.com/jacotton/LaVIISWA_genomes.

### Nucleotide diversity

Nucleotide diversity (pi, π) was computed from high-confidence bi-allelic filtered SNPs using vcftools 0.1.15 [70]. The genome-wide nucleotide diversity was calculated from a list of positions for each of the time points (pre- and post) and treatment groups (standard and intensive) using the option in vcftools ‘--site-pi’ respectively. The average nucleotide diversity within each of the groups was calculated individually and the symmetric 99% bootstrap confidence intervals of the averages were estimated using R version 3.5.1. Statistical significance of differences between group means was assessed by whether the confidence interval for one mean was disjoint from the mean of other groups. Effective population size (N_e_) was estimated from nucleotide diversity using the relationship π = 4.N_e_.µ [71] with the mutation rate 8.1 × 10^−9^ [57].

### Determination of rare allele sharing and kinship analysis

To identify the pairwise rare allele sharing we used a Perl script from Shortt et al. [47] available at https://github.com/PollockLaboratory/Schisto. We filtered for minor allele frequency ≤ 0.1 and sampled 500 SNP sites in 30 different generations. We then computed the mean value from the 30 generations for each pair. Allele-sharing scores were visualised in R version 4.0.2 using igraph v1.2.6 [72]. Significance of differences in mean allele sharing between groups were calculated against a non-parametric null distribution for each comparison generated by randomly permuting group labels 1000 times and calculating differences in mean allele sharing for each permutation.

### Test for selection

To test for recent positive selection within the treatment arms and between pre-treatment and post-treatment, the cross-population extended haplotype homozygosity (XP-EHH) test [73] was performed. XP-EHH is designed to detect whether either an ancestral or derived allele is undergoing selection within a given population. The XP-EHH test has the power to detect weaker signals of selection as it compares two closely related populations giving a directional score. The XP-EHH detects selective sweeps in which the selected allele has approached fixation in one population but remained polymorphic in another population. A VCF file containing only bi-allelic SNPs was subset into respective chromosomes. A genomic linkage map for each of the chromosomes was computed for each individual chromosome using the adjusted map length in centimorgan (cM) for the respective chromosomes [74]. The haplotypes from each of the chromosomes were then phased separately with their respective genomic map using Beagle v5.0 [75]. The XP-EHH test was performed using Selscan v1.2.0a [76] and the output XP-EHH scores were normalised for subsequent analysis using the norm program distributed with v1.2.0a of Selscan. Functional enrichment was assessed using g:Profiler version (e99_eg46_p14_f929183) [77] at a g:SCS threshold of 0.05 against a background of all annotated genes in *S. mansoni,* revealing genes showing significant purifying selection among the intensive and post-treatment parasite populations.

### Estimate of per-individual egg reduction rate (ERR) and association test

The posterior distribution for the ERR based on data from each individual for whom both pre- and post-treatment egg count data were available was estimated using a generalised linear mixed-effect model [78], incorporating nested random effects for treatment arm, village and individual. Means of the marginal posterior distribution per individual were used as quantitative phenotypes for an association study, testing all 6,967,554 called SNP variants. The model used was a linear regression of each SNP genotype against mean ERR, using 20 principal components as covariates to control for population structure, calculated using the ’--linear’ and ’--pca’ flags in plink v1.9 [79]. Code for these analyses is available at https://github.com/jacotton/LaVIISWA_genomes.

## Results

### Population stratification

After filtering, 6,967,554 high-confidence SNPs were retained (out of 18,716,072 unfiltered SNPs) from 174 individual miracidia. Principal components analysis of these high confidence SNPs showed little genetic structure within the island parasite populations on the first four principal components, which together represent 49% of the genetic variation. In particular, we found no evidence of population stratification between the standard (annual treatment) and intensive arms (quarterly treatment) or between pre- and post-treatment samples from these fishing communities (S2 Figure). The shoreline samples (from Mayuge district) clustered more closely with the island (Mukono district) parasite populations as compared to the inland samples (from Tororo district) (Fig. 2), but inland parasites were distinct from most island samples on principal component 3. The large-scale geographical pattern reflects the known genetic differentiation between inland and shoreline populations [55]. We also found that the island population is strikingly more diverse than either of the other populations (Fig. 2). While this is partly due to the larger number of samples included here, a larger sample of the shoreline and inland populations studied elsewhere also did not appear as diverse as the island population [55]. A number of miracidia appeared quite distinct from the main cluster of individuals on principal component 2. These divergent parasites were mostly (8 out of 9) from the islands and came from four different villages (Busi, Kakeeka, Zingoola, Katooke), with one from a shoreline village (Bwondha; S4 figure). Although participants were all resident in the villages throughout the LaVIISWA trial for at least 3 years before this study, there is a great deal of migration to the islands from other parts of the shoreline of Lake Victoria, including Kenya and Tanzania; therefore, we suspect these miracidia represent parasites imported from other populations that we have not sampled here.

**Fig 2.**
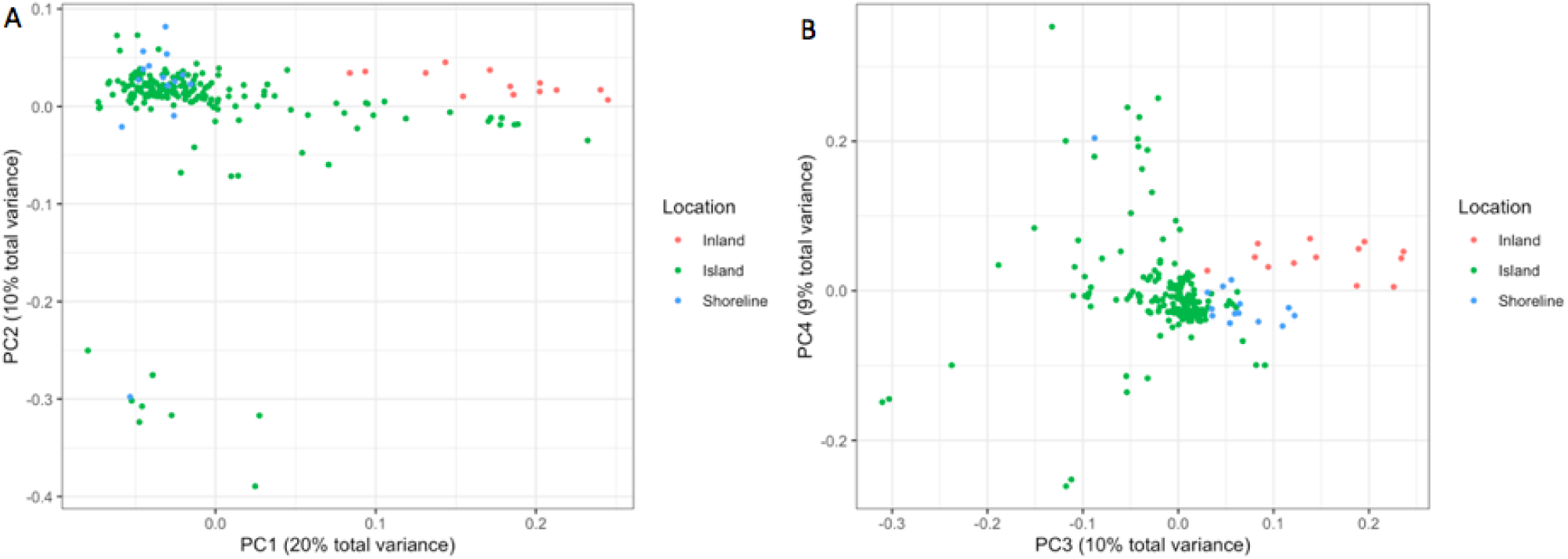
Principal components analysis of genetic variation within study samples and comparator Ugandan populations. (A) Shows the first two principal components and (B) the third and fourth principal components. Each point represents a single miracidium, coloured by the population from which they are sampled, with ’Shoreline’ samples from Mayuge district and ’Inland’ from Tororo district.

### Rare allele sharing and kinship analysis

To investigate direct relatedness between individual parasites, we adopted an approach based on determining the level of sharing of rare alleles (defined by their population frequency being less than or equal to 10%) between samples [47]. This approach has recently been used to study *S. japonicum* populations in China with whole-genome data [58]. By definition, most unrelated individuals share very few rare alleles; here we found slightly higher average proportion of rare-allele shared between pairs of miracidia isolated from the same individuals (0.1028) than in other comparisons (from the same village 0.0874, between villages on the same island 0.0862, between islands 0.0856). Differences between average allele sharing proportions were significant for comparing infrapopulation and within village groups (observed difference 0.0154, p-value from permutation test p<0.001) and within-village to within-island groups (observed difference 0.0012, p∼0.002) but only marginally so for within-island and between-island comparisons (observed difference 0.0006, p∼0.011). These data suggest an increase in relatedness within populations and possibly some geographical signature of increased relatedness.

We found three miracidia collected from the same infected individual at the same time with rare allele sharing of at least 0.3, and used these to calculate the average allele sharing for first-degree relatives (full siblings or parent-offspring) of 0.403 (actual values 0.4, 0.4014 and 0.409). This value is slightly lower than the theoretically expected level of identity-by-descent of 0.5, but with only three observations it is not possible to exclude that this difference is due to chance. Similarly, the average rare allele sharing for pairs of miracidia from different islands was 0.086, which represents our best estimate for the level of sharing in unrelated individuals. The small number of miracidia available for putative first-degree relatives means the observed variance from our data is very small, and so our final classification is thus deterministic. We classified miracidia pairs sharing more than 0.3105 of these rare alleles as first-degree relatives, 0.1981-0.3104 as second-degree relatives, 0.1419-0.1980 as third-degree relatives, 0.1138-0.1418 as fourth-degree relatives and 0.0956-0.1138 as fifth-degree relative, while those with less than 0.0956 sharing were classified as unrelated.

While 24% of pairs of samples from the same individual were classified as being related, only 10% of other comparisons appeared related (Fig. 3 A,B; 𝜒^2^ = 21.785, 1 df, p = 3.05 x 10^-6^), and a similar pattern held for close relatives (Fig. 3 A,B; first and second degree relatives represented 4% of within-infrapopulation comparisons, but 0.05% of all comparisons; 𝜒^2^ = 183.37, 1 df, p < 2.2 x 10^- 6^). There was no significant enrichment in related pairs of miracidia with either treatment intensity or for samples collected pre- and post-treatment. We found five pairs of first-degree relatives in total (Fig. 3C); but one pair were from different islands (marked 1 on Fig. 3C) and a second pair were from different individuals sampled on consecutive days in Kakeeka village (marked 2 on Fig. 3C). On the face of it, this would imply that the same combination of clonal cercariae infected these people, which seems very unlikely – particularly for the geographically separated cases. We cannot exclude the possibility that either the high level of rare allele sharing is misleading in these cases, or errors in sample identification. The remaining three pairs of first-degree relatives were pairs of miracidia sampled from single individuals in Busi, Kitosi and Kisu villages on consecutive days. Interestingly, a miracidium sampled from the same individual in Kisu was a second-degree relative of the first-degree pair, but this was collected post-treatment 37 days later. This is one of only 8 pairs of second-degree relatives. This suggested that either an adult worm survived treatment but changed ’partners’ (to produce a half-sibling or avuncular relationship) during this period, or two clonal worms with genetically distinct partners were present in this host at the two timepoints and produced these miracidia.

**Fig 3.**
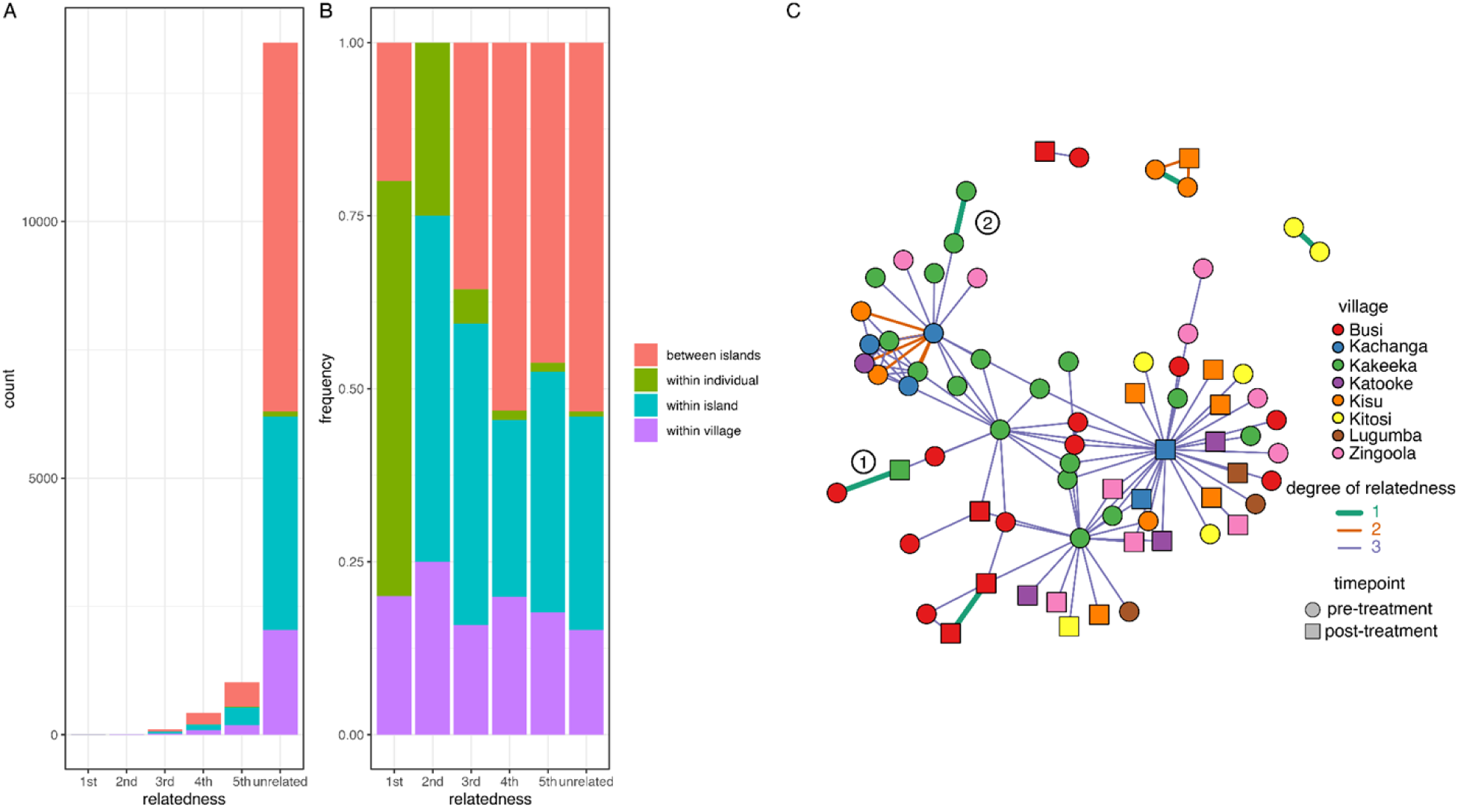
Patterns of relatedness inferred from pairwise rare allele sharing. (A) Number and (B) proportion of pairs of miracidia showing each degree of relatedness for miracidia sampled from the same individuals, villages or islands and for those on different islands. (C) Network representation of 1st, 2nd and 3rd degree relatedness. Vertices represent individual miracidia sampled, coloured by village and with a circle for samples taken pre-treatment and square for post-treatment samples. Edges join vertices inferred to share 1st, 2nd or 3rd degree relatedness, as indicated by both the width and colour of each edge. Numerical labels indicate two 1^st^ degree relationships discussed in the text.

### Analysis of genetic differentiation between villages

To further investigate genetic structure within the island population, we calculated F_ST_ (the proportion of genetic variation explained by population structure) for each pair of villages. As we expected, F_ST_ between villages was very low (maximum 0.0067), indicating little or no geographic structure to our data. We observed higher genetic differentiation between villages on different islands compared to those within the same island, but the small number of pairwise comparisons (N = 8 villages, 28 pairwise comparisons) meant that we did not have sufficient statistical power to detect any difference (p = 0.082, 1-way ANOVA of between/within village vs F_ST_). The villages were between 1 and 13 km apart, but there was no significant relationship between the distance between villages and F_ST_ (Fig. 4). To explore the geographical structure in these data more fully, we also fitted a gravity model attempting to explain F_ST_ between each pair of villages by the distance between villages, the population of each village and a factor capturing the effect of being on the same island. In this model, none of the explanatory variables had a significant influence on F_ST_, but the location of villages on the same island vs different islands was the most important variable with a likelihood weight in the best-fitting models of 0.48, while 0.31 for linear distance between villages and 0.23 for the product of village populations.

**Fig 4.**
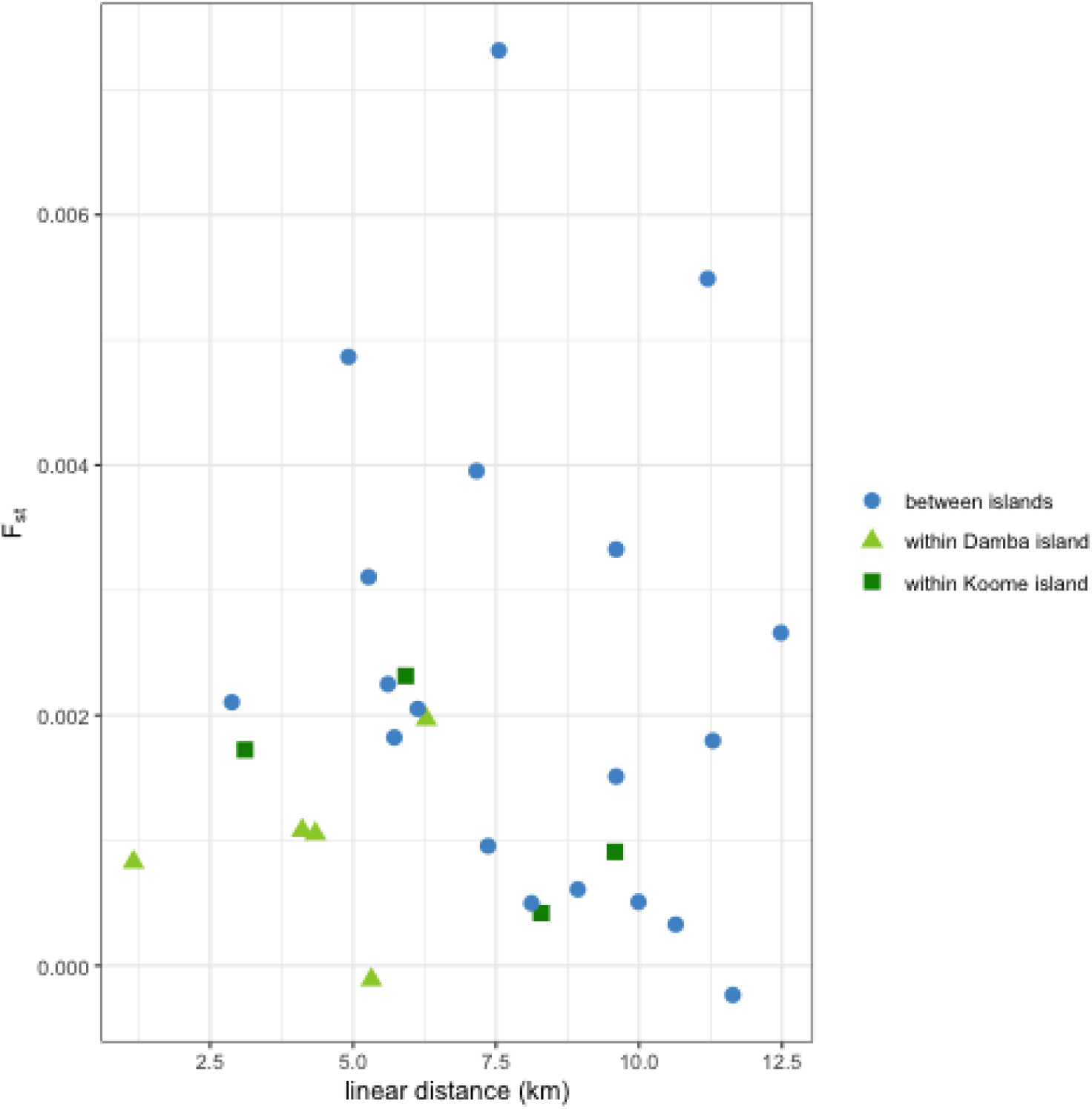
Pairwise F_ST_ estimates do not vary with linear distances between villages. Weir and Cockerham F_ST_ estimates used and distance measured in kilometres. Points show results of pairwise comparison between samples from different villages found on different islands or from different villages with Damba or Koome islands.

### Within-population genetic diversity

When comparing all pre- and post-treatment samples, we observed a very small but significant difference (p < 0.01) in genetic diversity between samples taken before and after treatment, with the 99% confidence interval for the mean nucleotide diversity in pre-treatment samples not overlapping with the mean post-treatment nucleotide diversity. This is consistent with a small effect of a single PZQ treatment round on the parasite population (Table 1). There was also lower diversity in parasites collected from villages in the intensive arm of the study than in the standard arm (Table 1), possibly reflecting a longer-term effect of more frequent PZQ treatment in these locations, despite the high levels of gene flow apparent between these locations implied by the very small levels of genetic differentiation we report. While this trend was consistent in both pre- and post-treatment samples, the difference between trial arms was most pronounced in post- treatment populations (Table 1). These diversity values are very similar to those observed in a recent study of the parasite populations on the lake shore and inland sites [55]. Using the mutation rate estimated previously [57], this implies an effective population size of around 10^5^ individuals from this sample collection, just outside the upper confidence limit of the estimate for the East Africa population in the previous study (3.67-9.35 x 10^4^) [57], and much higher than estimates from individual schools on the Lake Victoria shoreline (3.30-3.69 x 10^4^) [55], highlighting the diversity of *S. mansoni* parasites present on the islands.

**Table 1.**
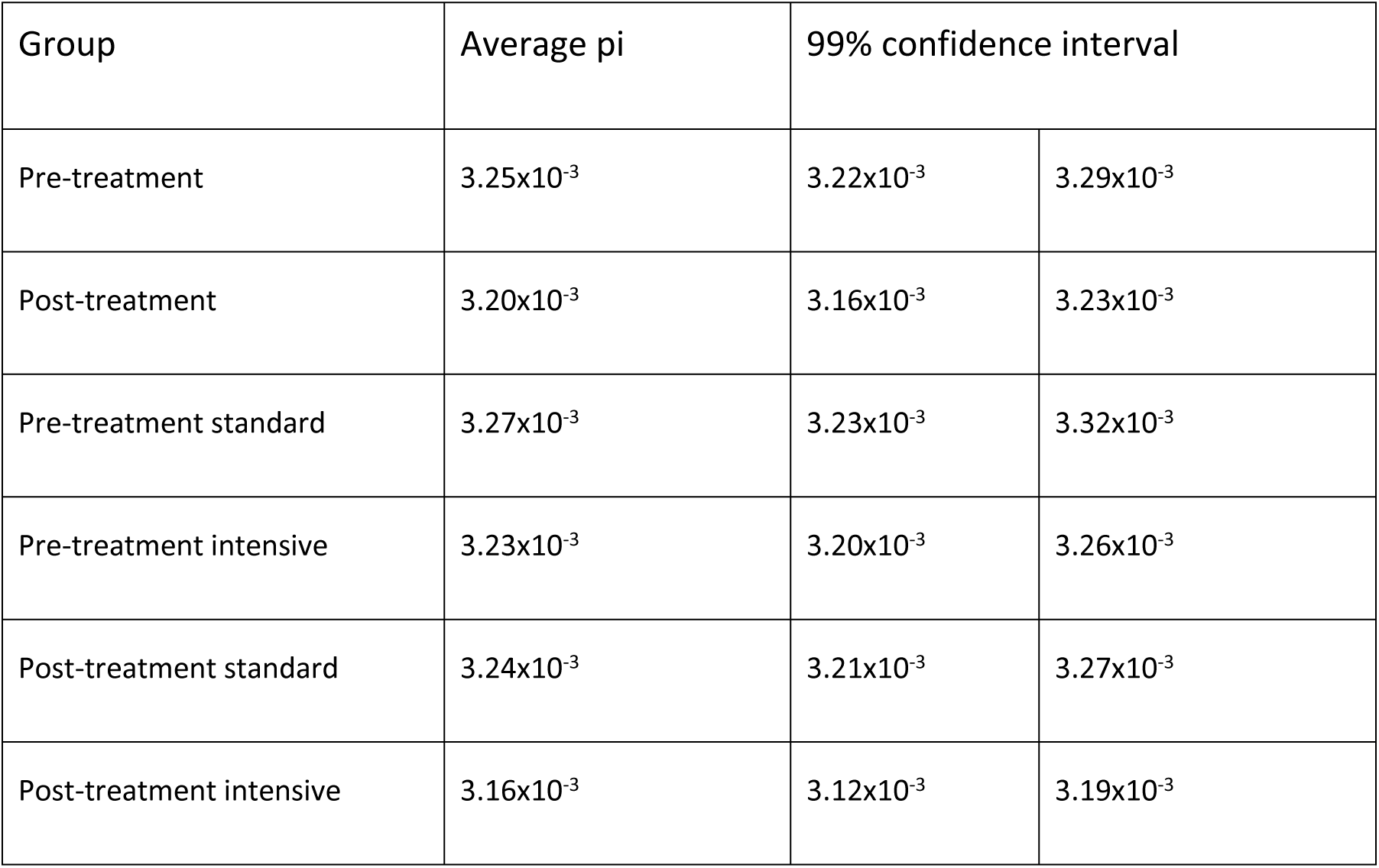
Genome-wide average nucleotide diversity (pi)

### Genetic differentiation with treatment between standard and intensive arms

Genome-wide average genetic differentiation was slightly higher (mean F_ST_ 3.9×10^-4^; bootstrap 99% CI: 2.5×10^-4^ - 5.3×10^-4^) between standard and intensive treatment populations post-treatment than before treatment (mean F_ST_ = 3.4×10^-4^; 99% CI = 1.8×10^-4^ – 5.0×10^-4^), but these values did not differ significantly. We also find very low genetic differentiation between standard and intensive trial arms (mean F_ST_ 5.6×10^-4^; 99% CI = 5.2×10^-4^ - 6.0×10^-4^ ). There was also variation in these F_ST_ values across the genome. While much of this likely reflects sampling variation (Fig. 5A), particularly striking was a region identified on chromosome 5 (Fig. 5B) with a distinct peak of divergence among post-treatment parasite populations. This window spanned 1.21 Mb of genomic sequence (from SM_V7_5: 7.78-8.99 Mb) and contained 25 annotated protein-coding genes (S2 table).

**Fig 5.**
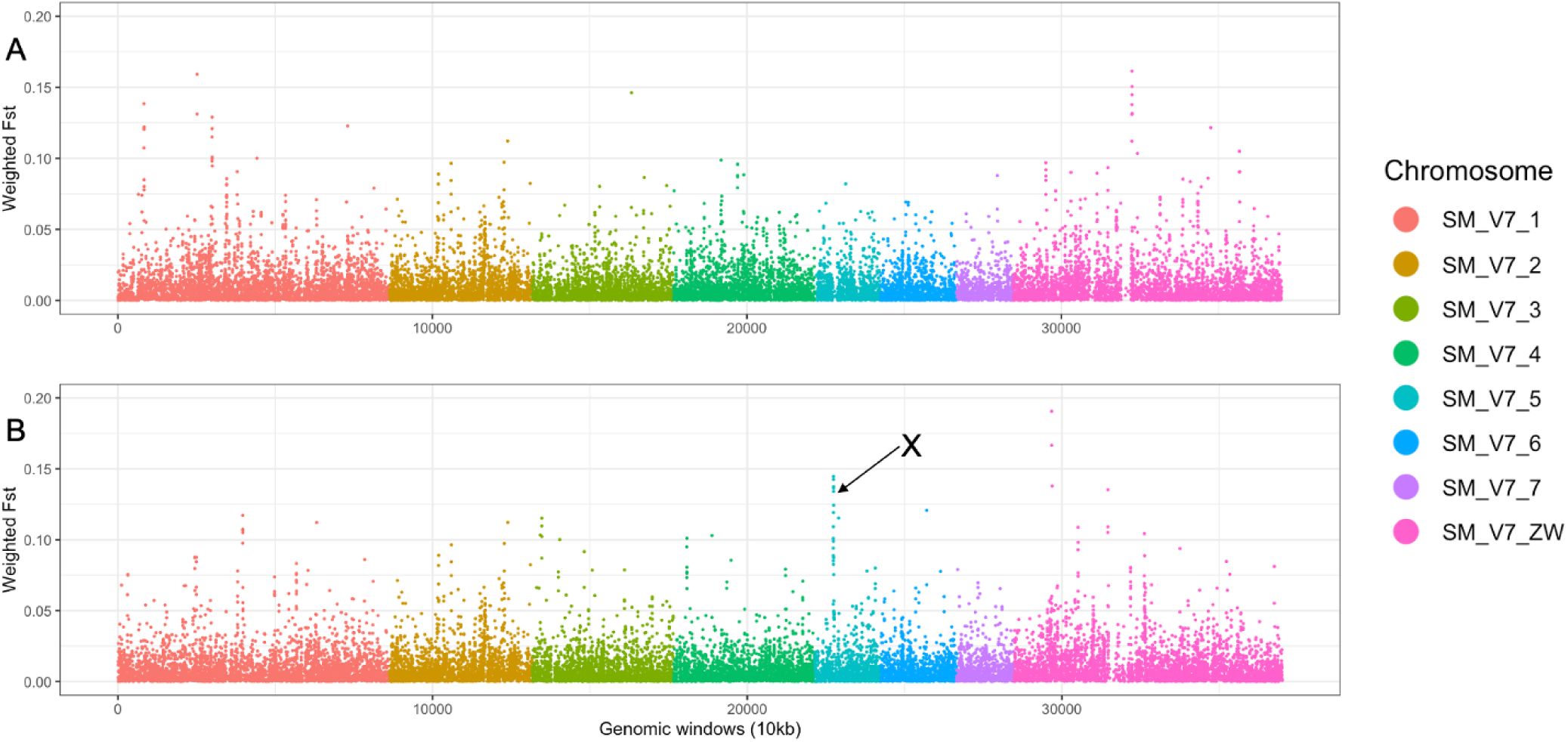
Genome-wide genetic differentiation between standard and intensive populations. F_ST_ calculated using pre-treatment (A) and post-treatment (B) samples. X marks the region of high post-treatment genetic differentiation discussed in the text. Each point represents the mean F_ST_ between genomic windows of 10 kb for all the called SNPs, with different coloured points representing SNPs on each chromosome.

### Analysis of signatures of selection

We used the XP-EHH test to identify genomic regions under differing selection pressures in separate comparisons between standard and intensive treatment arms (Fig. 6A) and between pre- and post-treatment samples (Fig. 6B). Taking extreme XP-EHH scores of < -2 or > 2 as a cutoff, we identified 510 windows as outliers including 12.75 Mb or 3.1% of the genome in total and representing 123 contiguous regions. None of the windows from either comparison overlap the peak of differentiation between standard and intensive treatment populations on chromosome 5. We note that the Z chromosome was particularly enriched for windows with extreme XP-EHH scores, containing almost half of those found genome-wide (5.325 Mb). This could be a technical artefact caused by difficulty in mapping to a highly repetitive chromosome [54], or due to the smaller average population size or a stronger effect of selection on recessive alleles when hemizygous. There are also a number of reasons to expect sex-linked genes to frequently be under selection [80]. An increased variance in XP-EHH scores is apparent specifically in the Z-specific region (Fig. 6) of the assembly scaffold representing the Z chromosome [54]. This region is not more repetitive than the autosomes or the pseudo-autosomal region shared by Z and W [see table S12 of 54], but is at lower copy number in the population as it is present in a single copy in female worms, so we suspect this enrichment of extreme XP-EHH scores represents a population genetic effect rather than a technical artefact.

**Fig 6.**
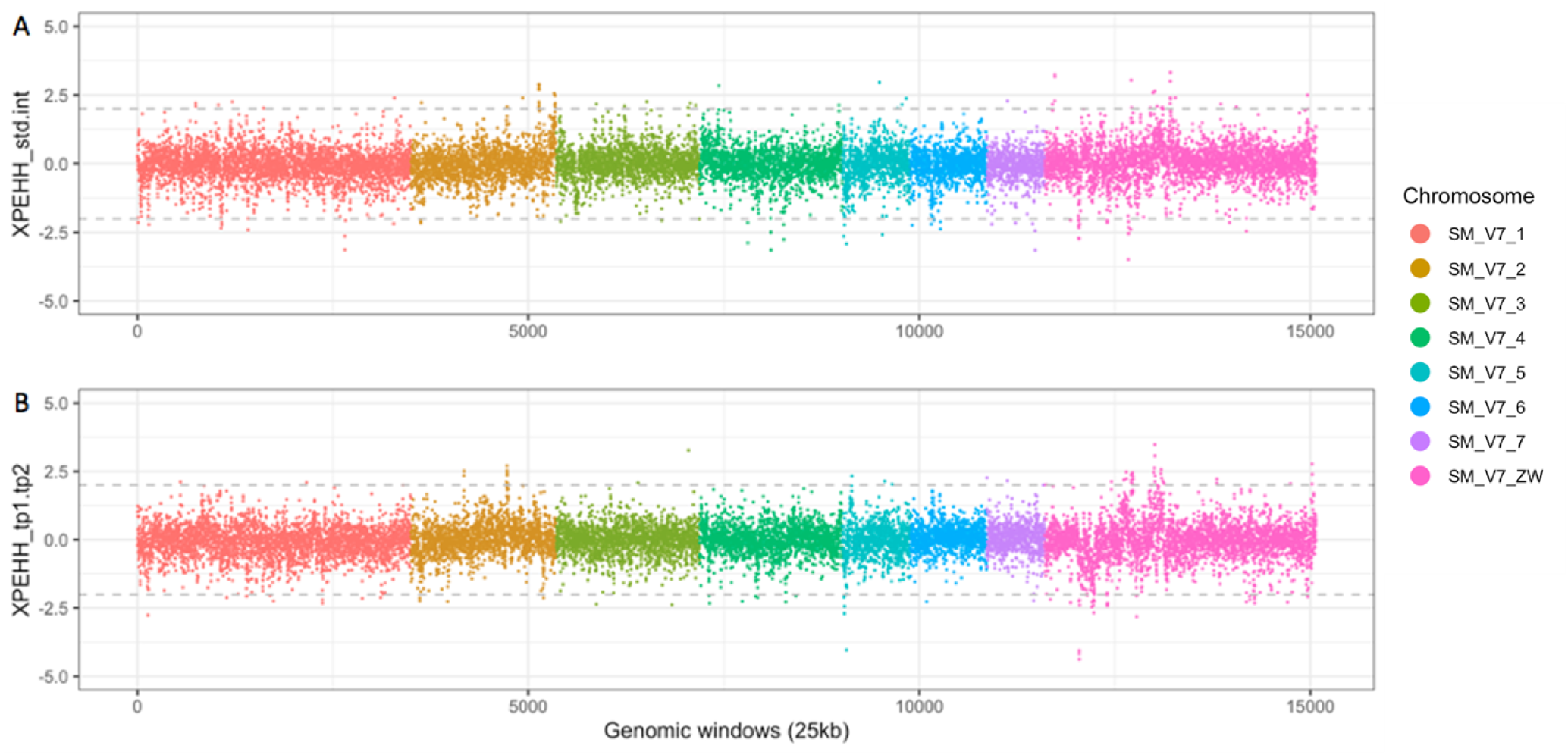
XP-EHH coloured by chromosome among treatment groups. A. Comparison between standard and intensive treatment groups. B. Comparison between pre-treatment and post-treatment groups. Positive values in panel A represent windows under stronger selective pressure in annual vs quarterly treatment arms. In panel B, positive values represent windows under stronger selection in pre-treatment than post-treatment samples.

There were 107 genes overlapping the outlier windows in the post-treatment samples, which were enriched for genes associated with seventeen GO terms for molecular function and biological processes (S3 table; https://biit.cs.ut.ee/gplink/l/wWE3Rp-ASq)55. Only 53 of these genes were on autosomes (leaving 54 on the Z and/or W chromosomes), and no GO terms were enriched when considering just the autosomal gene subset. No statistically significant enrichment for any functional category was observed among the genes undergoing stronger selection in pre-treatment individuals. Functional profiling showed that the 132 genes (78 autosomal) under stronger selection in the intensive arm were significantly enriched for association with 10 GO terms (S3 table; https://biit.cs.ut.ee/gplink/l/1VaAMWpxQK)55, which remained enriched in the autosomal subset. 88 genes were found in 46 autosomal windows with extreme XP-EHH values suggestive of stronger selection in the standard treatment arm include a pair of adjacent closely related genes likely to be a recent tandem duplication and possessing nucleoside deaminase activity on chromosome 4; these genes represent the only significantly enriched GO terms in this comparison (S3 table; https://biit.cs.ut.ee/gplink/l/4Fz7ZA3hTC).

### Individual egg reduction rate phenotypes

In an attempt to identify a phenotype for drug efficacy, we estimated the egg reduction rate (ERR) for 88 individuals for which genomic data was available and that had Kato-Katz egg counts taken both before and after treatment using a Bayesian linear mixed-effect model [78] that has previously been used to assess praziquantel efficacy [9, 81]. Previous analysis revealed a lower but not significant ERR in the intensive arm than the standard arm [10]. Similarly, we observe lower marginal ERR in samples collected in intensive than standard treatment villages, but with largely overlapping posterior distributions (Fig. 7B), and while villages vary in ERR (Fig. 7C), there were similar numbers of high- and low-clearance villages in the two arms. These differences were largely driven by a small number of individuals in some villages with very low (even negative - implying a higher egg count after treatment than before) ERR values (Fig. 7A). Unlike in a previous study [9], no ERRs were significantly below the 90% threshold, probably because only duplicate counts were available before and after treatment here, so there was significantly less information to estimate ERR on a per-individual basis.

**Fig 7.**
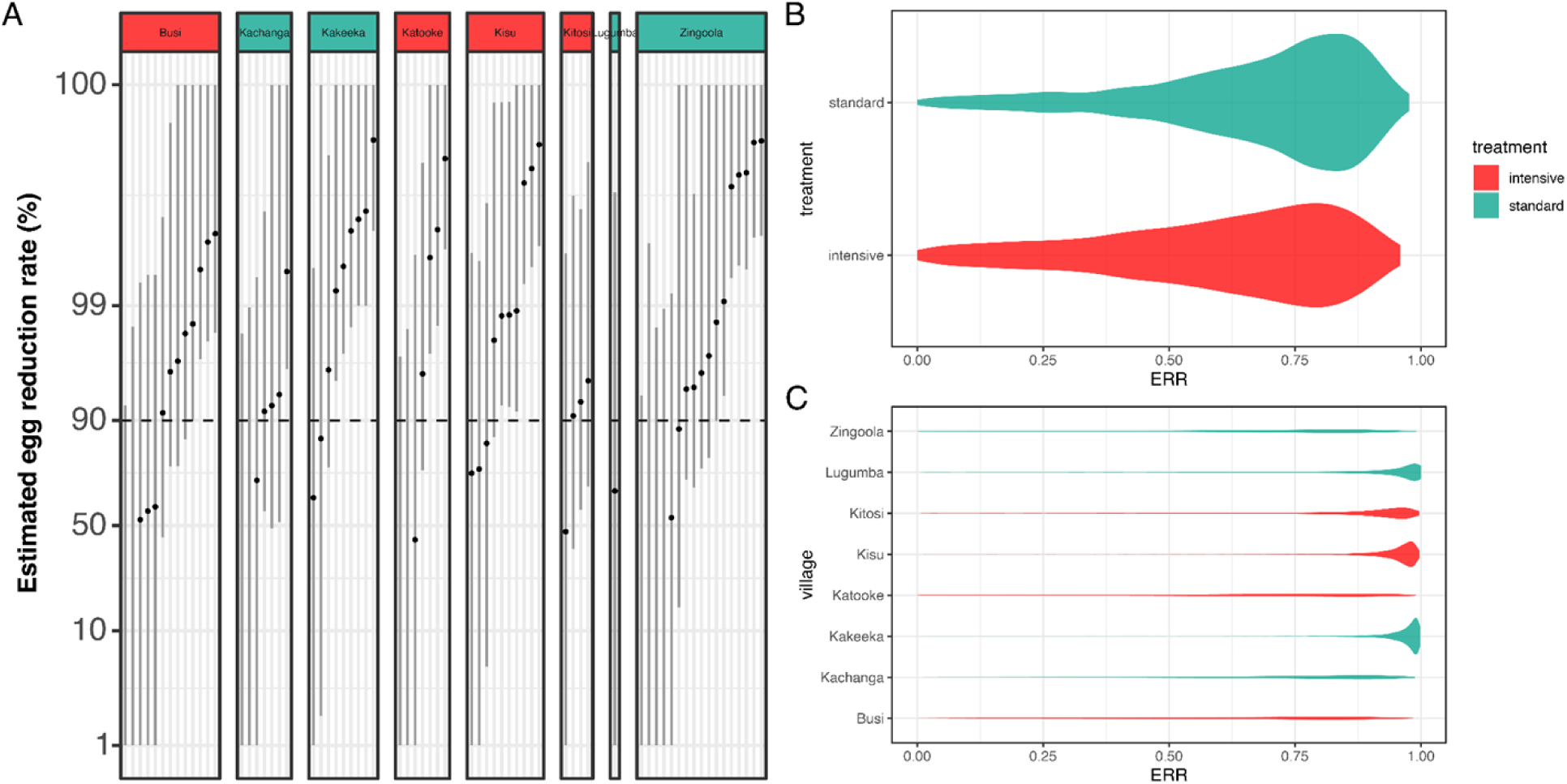
Egg reduction rate (ERR) estimates. (A) Posterior distributions of ERR for each individual for which pre- and post- egg count data were available. Lines indicate the 95% credible intervals (highest posterior density intervals) for each estimate, dots are the mean of the posterior distribution. Individuals are shown on an individual panel for each village, with panel headers coloured by treatment arm. Posterior distribution of average ERRs stratified by (B) treatment arm and (C) school were constructed by marginalizing over the fixed- and random-effects coefficients of the generalised linear mixed model.

Despite the small sample size, we attempted to identify genetic variants associated with differences in ERR, testing the 6.95 million high-quality SNPs found on the 7 autosomes or on the shared ZW scaffold. The smallest p-value for any SNP was 2.7×10^-9^, which after adjustment for multiple testing represents an adjusted p-value of 0.01841 (S3a Fig). There was some evidence that p-values are systematically biased in this analysis (S3b Fig). Correcting for population structure based on PCA coordinates removed the significance of hits (lowest p-value = 2.7×10^-8^; adjusted p-value = 0.179). The most significant hit (SM_V7_5:18325957) is intergenic, 886 bp upstream of an annotated protein-coding gene (Smp_314670) about which has no annotated domains or functional information are available. We thus conclude that there is no strong evidence linking any individual genetic variant in these data to variation in estimated ERR.

## Discussion

Schistosomiasis is second only to malaria in socio-economic impact among parasitic causes of morbidity and mortality [82–84]. MDA is the main method for schistosomiasis control, and there is currently an effort to expand the coverage of community-wide drug treatment to improve morbidity control [85] and address the persistence of schistosomiasis in some areas despite many years of PZQ distribution [86]. Understanding whether intensive treatment for individuals living in high transmission communities has an impact on parasite populations, potentially leading to drug resistance is of high importance for public health among schistosomiasis endemic communities in Africa. Determining the genetic basis of any drug resistance that does emerge is also crucial for tracking the spread of resistance through schistosome populations and for future drug or vaccine development designed to circumvent resistance as has been demonstrated for oxamniquine resistance [87].

Here, we have taken advantage of a large-scale trial in which the entire communities of 26 fishing villages were regularly treated with PZQ. A number of features of the study made this an ideal place to detect an effect of PZQ treatment on parasite populations. Villages were assigned randomly to treatment arms, so treatment frequency was independent of morbidity, parasite prevalence or intensity. Treatment was given under direct observation, avoiding issues with drug compliance reported in other studies [5]. The four week follow-up interval post-treatment would minimise the possibility of diagnosing newly acquired infection after treatment based on the development time of *S. mansoni* [88], as a new infection would take longer than four weeks to result in egg production that could be detected by Kato Katz and microscopy [89]. The exception would be if, during the time of treatment, a patient had juvenile worms as these would not have been cleared by treatment [90]. We expected that as the study was based on a group of islands it might help isolate the parasite population and so allow us to detect drug-induced selection in this population without the confounding effect of high levels of gene flow from untreated populations. Only individuals who had lived in these villages for at least three years were included in the study to control for absenteeism and MDA compliance although it was still not possible to control for movement between villages and islands given that fishing is the main economic activity within these communities.

The population of parasites present on the islands is closely related to that recently described from communities on the shoreline of Lake Victoria, and as expected rather divergent from that inland from the lake (Figs 2A & B) [55]. This presumably reflects greater movement of people between the shoreline and island than with the inland populations, as well as that the inland population included here is further (approximately 160 km) from the shoreline than are the islands (approximately 80 km). We see little genetic differentiation between villages on the same island, as fishing villages are close to one another (1-13 km apart) and movement may be frequent among fishermen and village communities. Less expected was that we see little or no genetic differentiation between islands, with only a weak trend for greater genetic differentiation between villages on different islands than villages on the same island, albeit this is a larger effect than either the distance between villages or the size of village populations, perhaps suggesting that snail vector movement around the coasts of islands may play a role in parasite movement. The islands are separated by water that is deep enough [91] (primary data at http://dataverse.harvard.edu/dataverse/LakeVicFish) to prevent snails moving actively from one island to the next, but parasites could travel through movement of infected people or through infected snails being carried on fishermen’s or conceivably by rafting on floating plants such as water hyacinth. Geographical conditions on these islands are similar except for Lugumba Island which has more rocky/stony shores compared to Koome and Damba which have more sandy shores and more vegetation, so we would expect snails to be able to establish similarly at most locations.

Despite seeing little or no genetic structure in the island parasite population, we see some evidence that PZQ treatment has had a small effect on the genetic diversity of the parasite population in this area. While we do not have baseline samples from before any PZQ treatment was administered as part of the LaVIISWA trial, we see very slightly higher genome-wide genetic diversity in the standard treatment arm than in the intensive arm, as would be expected if intensive treatment has been more effective at reducing the parasite population than the standard treatment regimen [10], although the effect we observe is very small and so maybe of limited biological relevance. Differences in the same direction were present when comparing subsets of samples taken before and after treatment separately, and was more pronounced in the post-treatment populations. As we see only very few closely related parasites, and no significant enrichment in relatedness based on treatment arm or sampling time with respect to treatment, it seems that this effect is unlikely to be due to differences in the number of directly related miracidia. We observe little or no genetic differentiation between villages in the two study arms, and only very slightly higher differentiation between the arms in post-treatment than in pre-treatment samples.

Evidence that PZQ treatment has some effect on the parasite population led us to investigate whether particular variants might be related to exposure to PZQ and so potentially responsible for any reduced susceptibility of parasites to PZQ within MDA programs [9]. We identify several regions within the genome that were highly differentiated between samples from the standard and intensive arms of the study, including a particularly striking region on chromosome 5 that showed high differentiation between post-treatment samples from the two arms of the study. This region contained a number of genes with functions that could be potentially linked to PZQ drug action. These include an ATP-binding cassette (ABC) transporter-associated gene (Smp_136310) that has previously been linked to helminth detoxification and drug resistance processes [92]. A gene with calcium dependent/modulatory functions (Smp_347070) was also found in the enriched region on chromosome 5, which is of interest given that the mode of action of PZQ has long been linked to increased permeability of the cell membrane to calcium ions into the cells which then causes contraction, paralysis and eventual death of the worms [93]. We also investigated regions of the genome under different selective regimes either with treatment intensity or when comparing pre- and post-treatment samples. Among the genes under varying selection were purine-nucleoside phosphorylase activity associated genes (Smp_197110 and Smp_171620) which are involved in the nucleotide salvage pathway of *S. mansoni*. Given that *S. mansoni* depends entirely on the salvage pathway for its purine metabolism [94], there is a possibility that ongoing non-random selection within this gene might affect parasite metabolic processes and a potential future drug target. However, we note that many biological processes could be contributing to genetic variation between samples from natural populations apart from variation in drug susceptibility [95]. While this study has shed some light on possible drug resistance genetic markers, other approaches, such as genetic crosses between parasites [96, 97] from natural populations that vary in drug efficacy or from lines selected for resistance [21, 87], are likely to have more power to reveal the genetics of drug resistance and so enable more focused studies of the effect on treatment on parasite populations.

A limitation in this study was that we did not have parasite populations sampled several years apart since it has been observed in similar studies that differentiation occurs over time in a given community [38], so sampling over a longer time-span could provide stronger evidence of genetic change in the population. In particular, we would ideally have access to baseline samples from the same population taken prior to any large-scale PZQ treatment being administered. Despite the falling cost and rising throughput of nucleic acid sequencing, we were limited in the number of miracidia that we could sequence in this study. An additional limitation is the labour-intensive process of hatching and washing miracidia necessary to obtain high-quality data due to the non-selective nature of the whole-genome sequencing approach [67].

As control programmes expand and reduce pathogen populations, we would expect the genetic diversity of these populations to fall to reflect the reduced population size [13, 41, 98], and drug resistance to be reflected in particular genotypes being over-represented in samples collected after large-scale treatment has been applied. As in other recent studies [42, 55], we find evidence of at best a very limited effect of PZQ treatment on schistosome populations either post-treatment or over a longer time frame of intensive treatment. In most previous studies, extensive refugia from treatment have been present in the community, as only school-age children are routinely treated in most areas, so it is instructive that we find similar results in this study despite community-wide treatment. While there is some evidence for reduced efficacy of PZQ in Uganda [9], most studies do not find a significant effect [11], including one study based on the same population as studied here [10]. Even in the absence of drug resistance emerging in natural populations, high-resolution genetic surveillance of African schistosome populations is ideally suited to detect changes in parasite population structure related to the impact of control measures [30], and could ultimately inform approaches to eliminate schistosome morbidity in remaining ’hot-spots’ by helping us understand parasite transmission between hosts and between foci [86].

In summary, we demonstrate a small but significant effect of both short-term PZQ treatment intensity and a recent treatment episode on genome wide-diversity in a schistosome. This reduction in diversity does not appear to be associated with enrichment of closely related parasites, but rather could reflect ongoing non-random recent selection within these fishing communities in Uganda that might be under the influence of continued mass drug administration. We identify genomic windows that are either particularly differentiated following treatment or appear to be under differing selective regimes with different treatment intensity. These regions could include genes involved in drug response, but additional data is needed to prioritise candidates for further investigation.

## Acknowledgements

We thank the Cure Rates study team and participants, the Wellcome Sanger Institute sequencing team, and members of the Parasite Genomics group and Pathogen Informatics team in the Parasites and Microbes Programme at the Sanger Institute. For the purpose of Open Access, the authors have applied a CC BY public copyright licence to any Author Accepted Manuscript version arising from this submission.

## Funding

This work was funded by the Wellcome Trust [grants 206194 and 095778].

## Conflict of interest

The authors declare that there are no conflicts of interest.

---

**S1 Figure.**
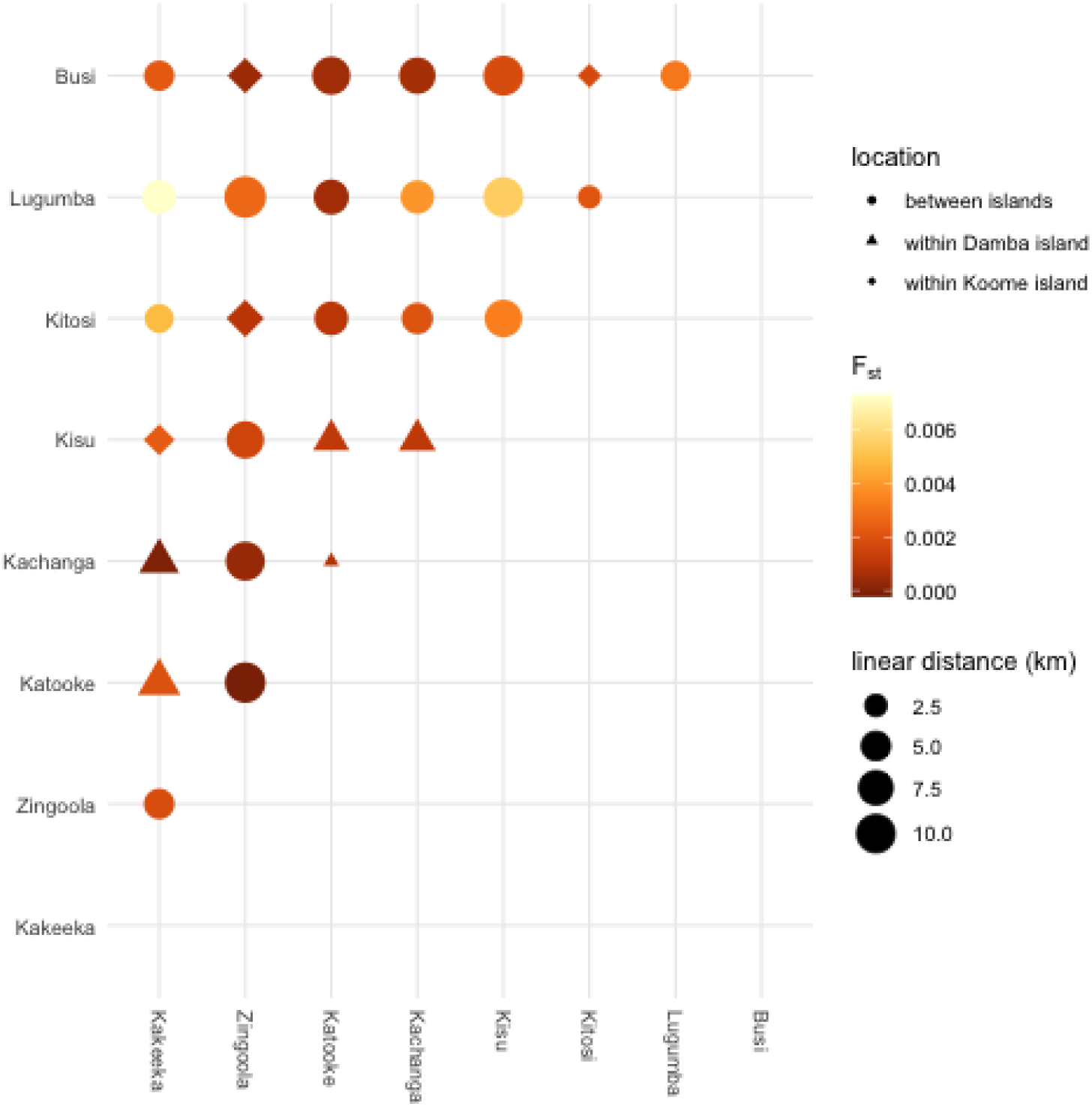
Mean F_ST_ between all pairs of villages. Shading indicates levels of genetic differentiation between pairs of villages indicated on each row and column. Symbol shapes reflect pair-wise comparisons of differentiation of populations samples between or within islands, and the area of each symbol is proportional to the linear distance between the villages.

**S2 Figure.**
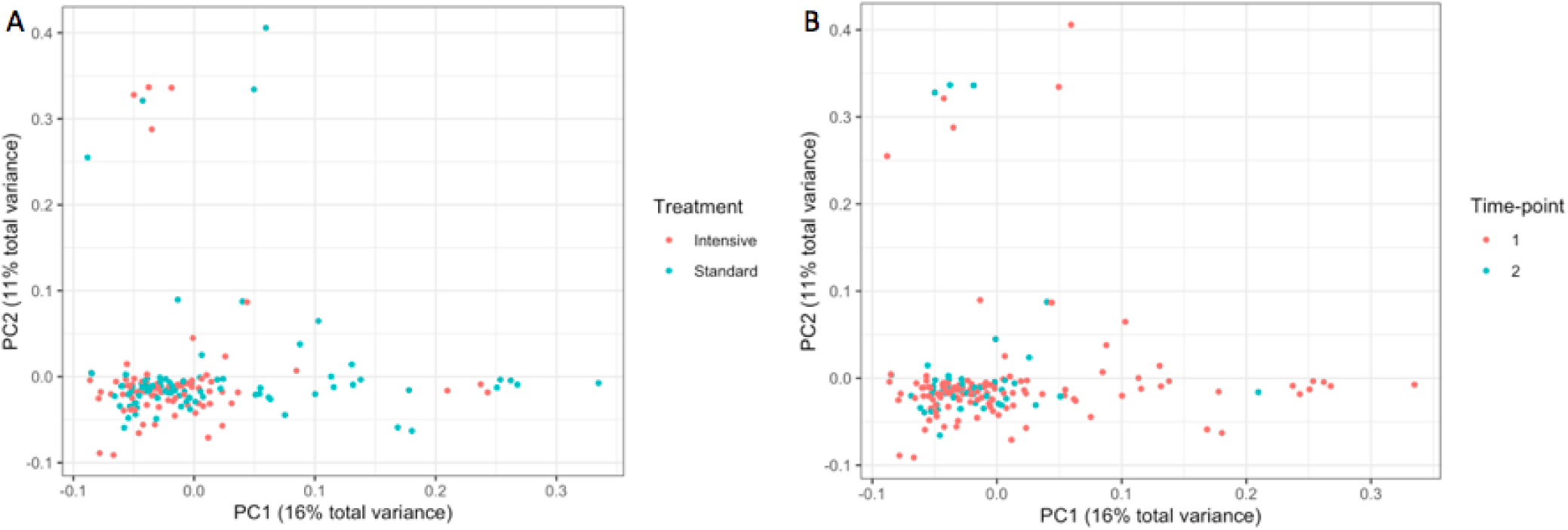
Principal component analysis of population structure by treatment and time point.

**S3 Figure.**
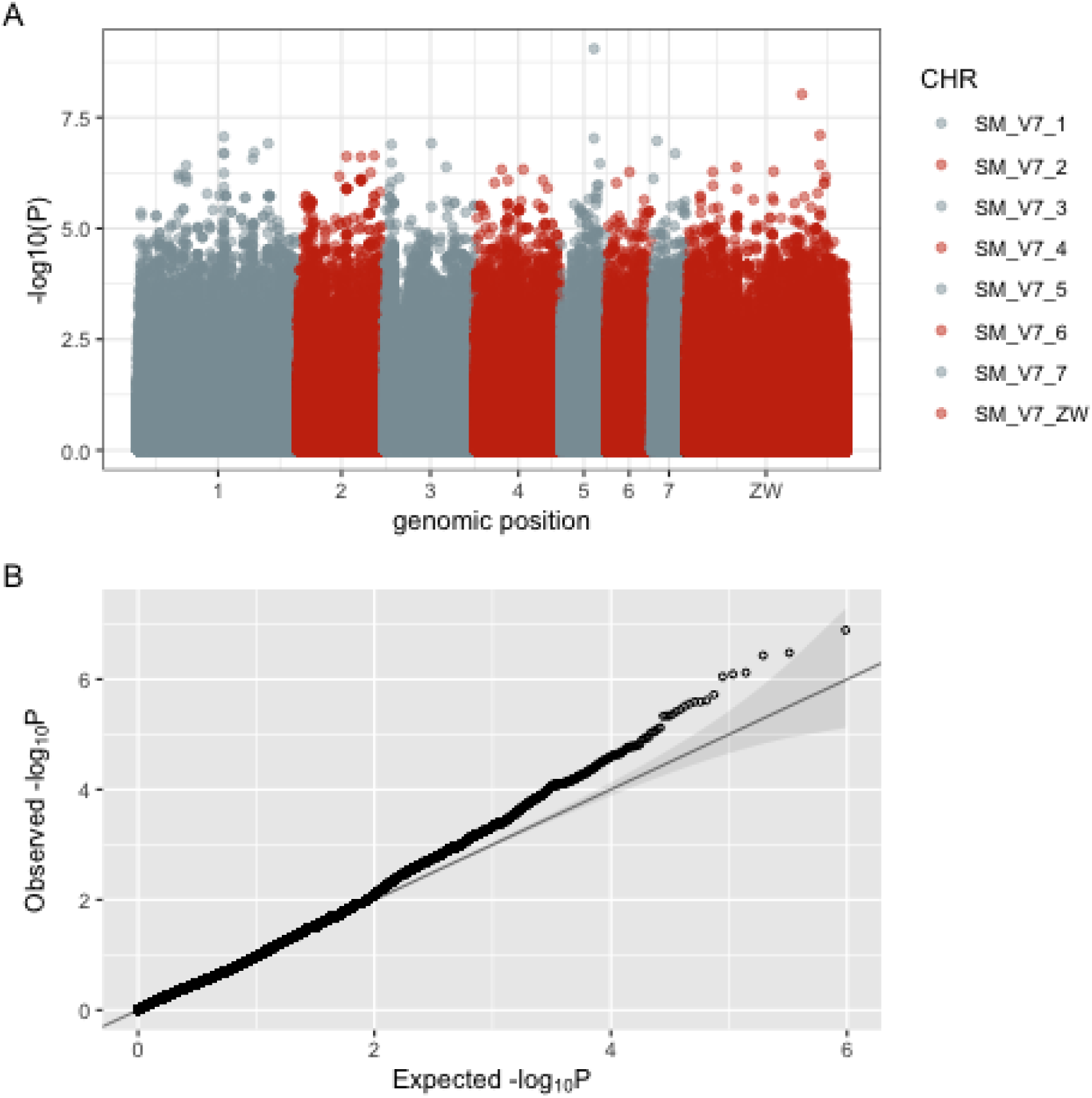
(A) Manhattan plot of unadjusted -log_10_ p-values for association of individual SNPs with per-individual mean egg-reduction rates. (B) QQplot of p-values from the same analysis against expectations under the null hypothesis.

**S4 Figure.**
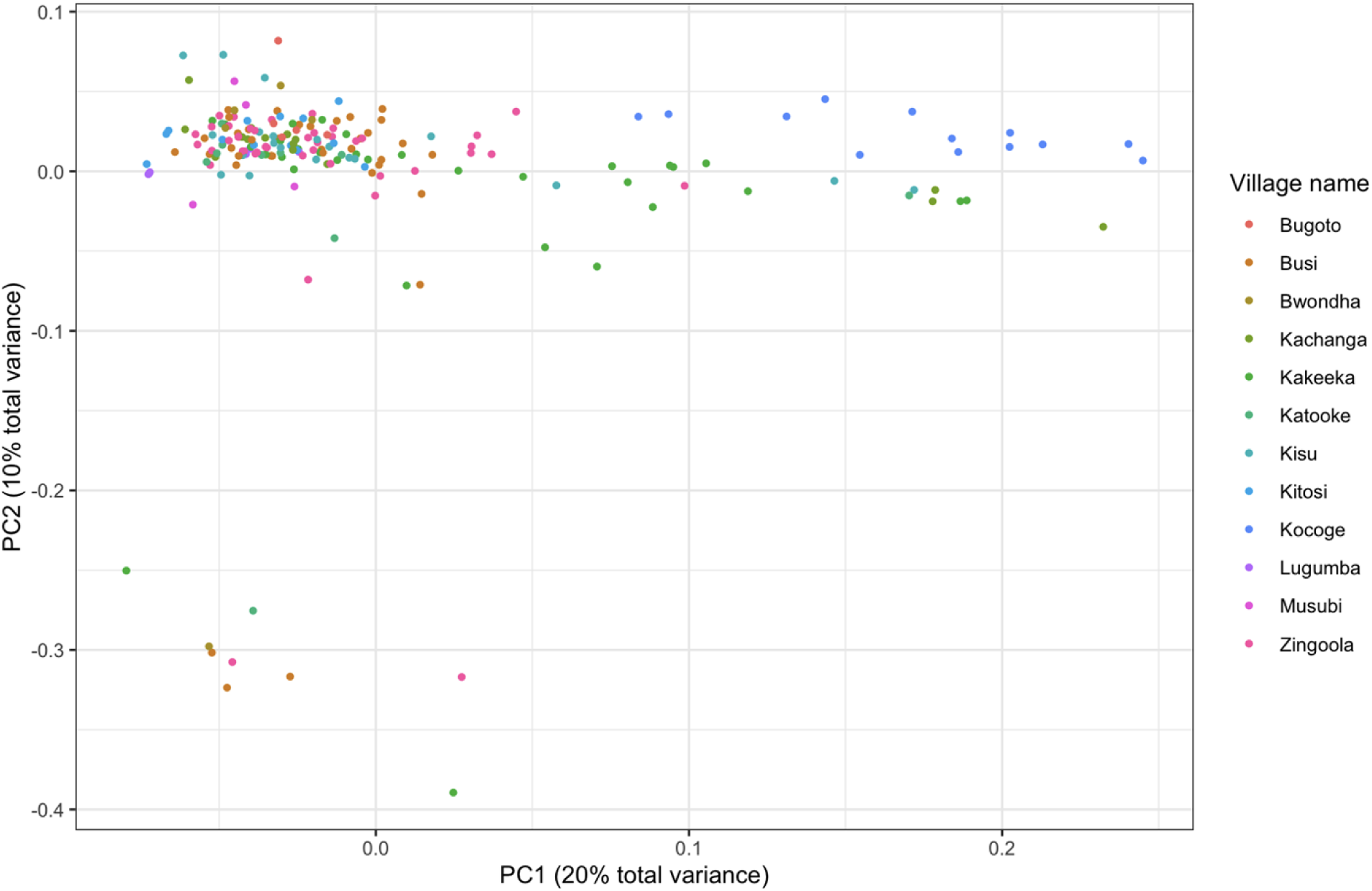
Principal component analysis showing a cluster of nine distinct miracidia on principal component 2 in the lower left quadrant.

**S1 Table.** Accession numbers and metadata for all samples included in analyses.

| sample_ID | collection_date | village | treatment_arm | island | accession_number | Pre/post |
| --- | --- | --- | --- | --- | --- | --- |
| 5582STDY7724293 | 10/08/2017 | Kachanga | Standard | Damba | ERS2891555 | pre_treatment |
| 5582STDY7759949 | 26/09/2017 | Zingoola | Standard | Koome | ERS2983612 | post_treatment |
| 5582STDY7724231 | 23/08/2017 | Zingoola | Standard | Koome | ERS2891493 | pre_treatment |
| 5582STDY7724309 | 10/08/2017 | Kachanga | Standard | Damba | ERS2891571 | pre_treatment |
| 5582STDY7724319 | 15/11/2017 | Kakeeka | Standard | Damba | ERS2891581 | post_treatment |
| 5582STDY7770933 | 10/07/2017 | Katooke | Intensive | Damba | ERS3016692 | pre_treatment |
| 5582STDY7770941 | 11/07/2017 | Katooke | Intensive | Damba | ERS3016700 | pre_treatment |
| 5582STDY7771009 | 10/07/2017 | Katooke | Intensive | Damba | ERS3016761 | pre_treatment |
| 5582STDY7770939 | 18/10/2017 | Busi | Intensive | Koome | ERS3016698 | pre_treatment |
| 5582STDY7724259 | 18/10/2017 | Busi | Intensive | Koome | ERS2891521 | pre_treatment |
| 5582STDY7724252 | 22/11/2017 | Busi | Intensive | Koome | ERS2891514 | post_treatment |
| 5582STDY7770956 | 17/10/2017 | Busi | Intensive | Koome | ERS3016715 | pre_treatment |
| 5582STDY7724233 | 18/10/2017 | Busi | Intensive | Koome | ERS2891495 | pre_treatment |
| 5582STDY7724277 | 23/11/2017 | Busi | Intensive | Koome | ERS2891539 | post_treatment |
| 5582STDY7724243 | 18/10/2017 | Busi | Intensive | Koome | ERS2891505 | pre_treatment |
| 5582STDY7724260 | 22/11/2017 | Busi | Intensive | Koome | ERS2891522 | post_treatment |
| 5582STDY7770991 | 19/10/2017 | Busi | Intensive | Koome | ERS3016764 | pre_treatment |
| 5582STDY7724269 | 21/11/2017 | Busi | Intensive | Koome | ERS2891531 | post_treatment |
| 5582STDY7770964 | 10/10/2017 | Kakeeka | Standard | Damba | ERS3016723 | pre_treatment |
| 5582STDY7771101 | 11/10/2017 | Kakeeka | Standard | Damba | ERS3016807 | pre_treatment |
| 5582STDY7724274 | 11/10/2017 | Kakeeka | Standard | Damba | ERS2891536 | pre_treatment |
| 5582STDY7724320 | 14/11/2017 | Kakeeka | Standard | Damba | ERS2891582 | post_treatment |
| 5582STDY7771003 | 11/10/2017 | Kakeeka | Standard | Damba | ERS3016755 | pre_treatment |
| 5582STDY7724265 | 10/10/2017 | Kakeeka | Standard | Damba | ERS2891527 | pre_treatment |
| 5582STDY7724281 | 09/10/2017 | Kakeeka | Standard | Damba | ERS2891543 | pre_treatment |
| 5582STDY7724240 | 13/11/2017 | Kakeeka | Standard | Damba | ERS2891502 | post_treatment |
| 5582STDY7771099 | 22/08/2017 | Zingoola | Standard | Koome | ERS3016805 | pre_treatment |
| 5582STDY7771043 | 23/08/2017 | Zingoola | Standard | Koome | ERS3016784 | pre_treatment |
| 5582STDY7724322 | 24/08/2017 | Zingoola | Standard | Koome | ERS2891584 | pre_treatment |
| 5582STDY7770915 | 23/08/2017 | Zingoola | Standard | Koome | ERS3016674 | pre_treatment |
| 5582STDY7724285 | 09/08/2017 | Kachanga | Standard | Damba | ERS2891547 | pre_treatment |
| 5582STDY7724288 | 07/11/2017 | Lugumba | Standard | Lugumba | ERS2891550 | pre_treatment |
| 5582STDY7724312 | 07/11/2017 | Lugumba | Standard | Lugumba | ERS2891574 | pre_treatment |
| 5582STDY7770937 | 13/09/2017 | Kachanga | Standard | Damba | ERS3016696 | post_treatment |
| 5582STDY7724250 | 09/10/2017 | Kakeeka | Standard | Damba | ERS2891512 | pre_treatment |
| 5582STDY7771004 | 11/09/2017 | Kachanga | Standard | Damba | ERS3016756 | post_treatment |
| 5582STDY7759911 | 24/10/2017 | Kitosi | Intensive | Koome | ERS2983579 | pre_treatment |
| 5582STDY7759950 | 23/10/2017 | Kitosi | Intensive | Koome | ERS2983613 | pre_treatment |
| 5582STDY7759951 | 22/11/2017 | Kitosi | Intensive | Koome | ERS2983614 | post_treatment |
| 5582STDY7759895 | 25/10/2017 | Kitosi | Intensive | Koome | ERS2983558 | pre_treatment |
| 5582STDY7759975 | 27/11/2017 | Kitosi | Intensive | Koome | ERS2983632 | post_treatment |
| 5582STDY7770926 | 10/07/2017 | Katooke | Intensive | Damba | ERS3016685 | pre_treatment |
| 5582STDY7759885 | 27/09/2017 | Zingoola | Standard | Koome | ERS2983563 | post_treatment |
| 5582STDY7770928 | 27/09/2017 | Zingoola | Standard | Koome | ERS3016687 | post_treatment |
| 5582STDY7770942 | 09/10/2017 | Kakeeka | Standard | Damba | ERS3016701 | pre_treatment |
| 5582STDY7770948 | 11/10/2017 | Kakeeka | Standard | Damba | ERS3016707 | pre_treatment |
| 5582STDY7771093 | 10/10/2017 | Kakeeka | Standard | Damba | ERS3016804 | pre_treatment |
| 5582STDY7724242 | 09/10/2017 | Kakeeka | Standard | Damba | ERS2891504 | pre_treatment |
| 5582STDY7724297 | 11/10/2017 | Kakeeka | Standard | Damba | ERS2891559 | pre_treatment |
| 5582STDY7771022 | 10/10/2017 | Kakeeka | Standard | Damba | ERS3016777 | pre_treatment |
| 5582STDY7770987 | 12/10/2017 | Kakeeka | Standard | Damba | ERS3016746 | pre_treatment |
| 5582STDY7771035 | 23/08/2017 | Zingoola | Standard | Koome | ERS3016781 | pre_treatment |
| 5582STDY7759963 | 31/10/2017 | Kisu | Intensive | Damba | ERS2983622 | pre_treatment |
| 5582STDY7770938 | 06/12/2017 | Kisu | Intensive | Damba | ERS3016697 | post_treatment |
| 5582STDY7759939 | 31/10/2017 | Kisu | Intensive | Damba | ERS2983603 | pre_treatment |
| 5582STDY7759930 | 05/12/2017 | Kisu | Intensive | Damba | ERS2983595 | post_treatment |
| 5582STDY7759954 | 06/12/2017 | Kisu | Intensive | Damba | ERS2983615 | post_treatment |
| 5582STDY7770954 | 06/12/2017 | Kisu | Intensive | Damba | ERS3016713 | post_treatment |
| 5582STDY7770961 | 05/12/2017 | Kisu | Intensive | Damba | ERS3016720 | post_treatment |
| 5582STDY7759962 | 06/12/2017 | Kisu | Intensive | Damba | ERS2983621 | post_treatment |
| 5582STDY7770943 | 24/08/2017 | Zingoola | Standard | Koome | ERS3016702 | pre_treatment |
| 5582STDY7770986 | 24/08/2017 | Zingoola | Standard | Koome | ERS3016745 | pre_treatment |
| 5582STDY7759964 | 22/08/2017 | Zingoola | Standard | Koome | ERS2983623 | pre_treatment |
| 5582STDY7759965 | 26/09/2017 | Zingoola | Standard | Koome | ERS2983624 | post_treatment |
| 5582STDY7770951 | 24/08/2017 | Zingoola | Standard | Koome | ERS3016710 | pre_treatment |
| 5582STDY7770978 | 24/08/2017 | Zingoola | Standard | Koome | ERS3016737 | pre_treatment |
| 5582STDY7771067 | 22/08/2017 | Zingoola | Standard | Koome | ERS3016793 | pre_treatment |
| 5582STDY7759887 | 25/10/2017 | Kitosi | Intensive | Koome | ERS2983565 | pre_treatment |
| 5582STDY7759966 | 26/10/2017 | Kitosi | Intensive | Koome | ERS2983625 | pre_treatment |
| 5582STDY7759919 | 28/11/2017 | Kitosi | Intensive | Koome | ERS2983586 | post_treatment |
| 5582STDY7771000 | 19/10/2017 | Busi | Intensive | Koome | ERS3016752 | pre_treatment |
| 5582STDY7724238 | 08/08/2017 | Kachanga | Standard | Damba | ERS2891500 | pre_treatment |
| 5582STDY7724230 | 12/09/2017 | Kachanga | Standard | Damba | ERS2891492 | post_treatment |
| 5582STDY7724241 | 18/10/2017 | Busi | Intensive | Koome | ERS2891503 | pre_treatment |
| 5582STDY7724316 | 17/10/2017 | Busi | Intensive | Koome | ERS2891578 | pre_treatment |
| 5582STDY7771044 | 24/08/2017 | Zingoola | Standard | Koome | ERS3016785 | pre_treatment |
| 5582STDY7771075 | 22/08/2017 | Zingoola | Standard | Koome | ERS3016796 | pre_treatment |
| 5582STDY7724294 | 23/08/2017 | Zingoola | Standard | Koome | ERS2891556 | pre_treatment |
| 5582STDY7771002 | 24/08/2017 | Zingoola | Standard | Koome | ERS3016754 | pre_treatment |
| 5582STDY7759974 | 25/10/2017 | Kitosi | Intensive | Koome | ERS2983631 | pre_treatment |
| 5582STDY7770971 | 11/10/2017 | Kakeeka | Standard | Damba | ERS3016730 | pre_treatment |
| 5582STDY7724249 | 10/10/2017 | Kakeeka | Standard | Damba | ERS2891511 | pre_treatment |
| 5582STDY7770924 | 12/10/2017 | Kakeeka | Standard | Damba | ERS3016683 | pre_treatment |
| 5582STDY7724266 | 11/10/2017 | Kakeeka | Standard | Damba | ERS2891528 | pre_treatment |
| 5582STDY7724247 | 12/10/2017 | Kakeeka | Standard | Damba | ERS2891509 | pre_treatment |
| 5582STDY7724248 | 15/11/2017 | Kakeeka | Standard | Damba | ERS2891510 | post_treatment |
| 5582STDY7771014 | 10/10/2017 | Kakeeka | Standard | Damba | ERS3016773 | pre_treatment |
| 5582STDY7724301 | 10/08/2017 | Kachanga | Standard | Damba | ERS2891563 | pre_treatment |
| 5582STDY7771076 | 24/10/2017 | Kitosi | Intensive | Koome | ERS3016797 | pre_treatment |
| 5582STDY7770977 | 15/07/2017 | Katooke | Intensive | Damba | ERS3016736 | pre_treatment |
| 5582STDY7771011 | 23/08/2017 | Zingoola | Standard | Koome | ERS3016770 | pre_treatment |
| 5582STDY7770919 | 23/08/2017 | Zingoola | Standard | Koome | ERS3016678 | pre_treatment |
| 5582STDY7771091 | 22/08/2017 | Zingoola | Standard | Koome | ERS3016802 | pre_treatment |
| 5582STDY7724310 | 23/08/2017 | Zingoola | Standard | Koome | ERS2891572 | pre_treatment |
| 5582STDY7724272 | 07/11/2017 | Lugumba | Standard | Lugumba | ERS2891534 | pre_treatment |
| 5582STDY7724264 | 13/12/2017 | Lugumba | Standard | Lugumba | ERS2891526 | post_treatment |
| 5582STDY7770969 | 12/07/2017 | Katooke | Intensive | Damba | ERS3016728 | pre_treatment |
| 5582STDY7770979 | 12/10/2017 | Kakeeka | Standard | Damba | ERS3016738 | pre_treatment |
| 5582STDY7759883 | 06/12/2017 | Kisu | Intensive | Damba | ERS2983561 | post_treatment |
| 5582STDY7759900 | 30/10/2017 | Kisu | Intensive | Damba | ERS2983569 | pre_treatment |
| 5582STDY7759955 | 01/11/2017 | Kisu | Intensive | Damba | ERS2983616 | pre_treatment |
| 5582STDY7770922 | 06/12/2017 | Kisu | Intensive | Damba | ERS3016681 | post_treatment |
| 5582STDY7759904 | 17/08/2017 | Katooke | Intensive | Damba | ERS2983573 | post_treatment |
| 5582STDY7759916 | 01/11/2017 | Kisu | Intensive | Damba | ERS2983583 | pre_treatment |
| 5582STDY7724287 | 09/10/2017 | Kakeeka | Standard | Damba | ERS2891549 | pre_treatment |
| 5582STDY7724273 | 10/10/2017 | Kakeeka | Standard | Damba | ERS2891535 | pre_treatment |
| 5582STDY7724236 | 23/11/2017 | Busi | Intensive | Koome | ERS2891498 | post_treatment |
| 5582STDY7724244 | 22/11/2017 | Busi | Intensive | Koome | ERS2891506 | post_treatment |
| 5582STDY7724253 | 18/10/2017 | Busi | Intensive | Koome | ERS2891515 | pre_treatment |
| 5582STDY7724291 | 17/10/2017 | Busi | Intensive | Koome | ERS2891553 | pre_treatment |
| 5582STDY7770990 | 23/11/2017 | Busi | Intensive | Koome | ERS3016763 | post_treatment |
| 5582STDY7724315 | 23/11/2017 | Busi | Intensive | Koome | ERS2891577 | post_treatment |
| 5582STDY7771068 | 23/10/2017 | Kitosi | Intensive | Koome | ERS3016794 | pre_treatment |
| 5582STDY7724278 | 22/08/2017 | Zingoola | Standard | Koome | ERS2891540 | pre_treatment |
| 5582STDY7759941 | 27/09/2017 | Zingoola | Standard | Koome | ERS2983605 | post_treatment |
| 5582STDY7771036 | 24/08/2017 | Zingoola | Standard | Koome | ERS3016782 | pre_treatment |
| 5582STDY7771051 | 23/08/2017 | Zingoola | Standard | Koome | ERS3016787 | pre_treatment |
| 5582STDY7771007 | 18/10/2017 | Busi | Intensive | Koome | ERS3016759 | pre_treatment |
| 5582STDY7724308 | 18/10/2017 | Busi | Intensive | Koome | ERS2891570 | pre_treatment |
| 5582STDY7724276 | 22/11/2017 | Busi | Intensive | Koome | ERS2891538 | post_treatment |
| 5582STDY7759925 | 27/09/2017 | Zingoola | Standard | Koome | ERS2983591 | post_treatment |
| 5582STDY7759918 | 23/10/2017 | Kitosi | Intensive | Koome | ERS2983585 | pre_treatment |
| 5582STDY7759926 | 24/10/2017 | Kitosi | Intensive | Koome | ERS2983592 | pre_treatment |
| 5582STDY7724229 | 18/10/2017 | Busi | Intensive | Koome | ERS2891491 | pre_treatment |
| 5582STDY7724227 | 17/10/2017 | Busi | Intensive | Koome | ERS2891489 | pre_treatment |
| 5582STDY7724235 | 19/10/2017 | Busi | Intensive | Koome | ERS2891497 | pre_treatment |
| 5582STDY7724268 | 22/11/2017 | Busi | Intensive | Koome | ERS2891530 | post_treatment |
| 5582STDY7724300 | 23/11/2017 | Busi | Intensive | Koome | ERS2891562 | post_treatment |
| 5582STDY7770959 | 24/08/2017 | Zingoola | Standard | Koome | ERS3016718 | pre_treatment |
| 5582STDY7771059 | 22/08/2017 | Zingoola | Standard | Koome | ERS3016790 | pre_treatment |
| 5582STDY7771010 | 24/08/2017 | Zingoola | Standard | Koome | ERS3016769 | pre_treatment |
| 5582STDY7770929 | 27/09/2017 | Zingoola | Standard | Koome | ERS3016688 | post_treatment |
| 5582STDY7770935 | 24/08/2017 | Zingoola | Standard | Koome | ERS3016694 | pre_treatment |
| 5582STDY7724302 | 23/08/2017 | Zingoola | Standard | Koome | ERS2891564 | pre_treatment |
| 5582STDY7771028 | 24/08/2017 | Zingoola | Standard | Koome | ERS3016779 | pre_treatment |
| 5582STDY7759888 | 13/07/2017 | Katooke | Intensive | Damba | ERS2983566 | pre_treatment |
| 5582STDY7759928 | 16/08/2017 | Katooke | Intensive | Damba | ERS2983594 | post_treatment |
| 5582STDY7770944 | 13/09/2017 | Kachanga | Standard | Damba | ERS3016703 | post_treatment |
| 5582STDY7759936 | 15/08/2017 | Katooke | Intensive | Damba | ERS2983601 | pre_treatment |
| 5582STDY7759944 | 13/09/2017 | Kachanga | Standard | Damba | ERS2983608 | post_treatment |
| 5582STDY7770976 | 18/10/2017 | Busi | Intensive | Koome | ERS3016735 | pre_treatment |
| 5582STDY7770953 | 24/08/2017 | Zingoola | Standard | Koome | ERS3016712 | pre_treatment |
| 5582STDY7759894 | 28/09/2017 | Zingoola | Standard | Koome | ERS2983557 | post_treatment |
| 5582STDY7759901 | 28/09/2017 | Zingoola | Standard | Koome | ERS2983570 | post_treatment |
| 5582STDY7771008 | 17/10/2017 | Busi | Intensive | Koome | ERS3016760 | pre_treatment |
| 5582STDY7770982 | 07/08/2017 | Kachanga | Standard | Damba | ERS3016741 | pre_treatment |
| 5582STDY7770936 | 13/09/2017 | Kachanga | Standard | Damba | ERS3016695 | post_treatment |
| 5582STDY7770927 | 24/08/2017 | Zingoola | Standard | Koome | ERS3016686 | pre_treatment |
| 5582STDY7759933 | 26/09/2017 | Zingoola | Standard | Koome | ERS2983598 | post_treatment |
| 5582STDY7724286 | 23/08/2017 | Zingoola | Standard | Koome | ERS2891548 | pre_treatment |
| 5582STDY7770945 | 27/09/2017 | Zingoola | Standard | Koome | ERS3016704 | post_treatment |
| 5582STDY7770966 | 22/11/2017 | Busi | Intensive | Koome | ERS3016725 | post_treatment |
| 5582STDY7724245 | 19/10/2017 | Busi | Intensive | Koome | ERS2891507 | pre_treatment |
| 5582STDY7724313 | 18/10/2017 | Busi | Intensive | Koome | ERS2891575 | pre_treatment |
| 5582STDY7724237 | 18/10/2017 | Busi | Intensive | Koome | ERS2891499 | pre_treatment |
| 5582STDY7724304 | 08/11/2017 | Lugumba | Standard | Lugumba | ERS2891566 | pre_treatment |
| 5582STDY7770983 | 17/10/2017 | Busi | Intensive | Koome | ERS3016742 | pre_treatment |
| 5582STDY7759938 | 05/12/2017 | Kisu | Intensive | Damba | ERS2983602 | post_treatment |
| 5582STDY7771053 | 01/11/2017 | Kisu | Intensive | Damba | ERS3016789 | pre_treatment |
| 5582STDY7771061 | 31/10/2017 | Kisu | Intensive | Damba | ERS3016792 | pre_treatment |
| 5582STDY7759906 | 31/10/2017 | Kisu | Intensive | Damba | ERS2983574 | pre_treatment |
| 5582STDY7759898 | 07/12/2017 | Kisu | Intensive | Damba | ERS2983567 | post_treatment |
| 5582STDY7759882 | 06/12/2017 | Kisu | Intensive | Damba | ERS2983560 | post_treatment |
| 5582STDY7724317 | 13/09/2017 | Kachanga | Standard | Damba | ERS2891579 | post_treatment |
| 5582STDY7724263 | 11/10/2017 | Kakeeka | Standard | Damba | ERS2891525 | pre_treatment |
| 5582STDY7724279 | 10/10/2017 | Kakeeka | Standard | Damba | ERS2891541 | pre_treatment |
| 5582STDY7770916 | 11/10/2017 | Kakeeka | Standard | Damba | ERS3016675 | pre_treatment |
| 5582STDY7724271 | 09/10/2017 | Kakeeka | Standard | Damba | ERS2891533 | pre_treatment |
| 5582STDY7770957 | 07/08/2017 | Kachanga | Standard | Damba | ERS3016716 | pre_treatment |
| 5582STDY7770973 | 09/08/2017 | Kachanga | Standard | Damba | ERS3016732 | pre_treatment |
| 5582STDY7770950 | 10/10/2017 | Kakeeka | Standard | Damba | ERS3016709 | pre_treatment |
| 5582STDY7771029 | 01/11/2017 | Kisu | Intensive | Damba | ERS3016780 | pre_treatment |
| 5582STDY7770975 | 17/10/2017 | Busi | Intensive | Koome | ERS3016734 | pre_treatment |
| 5582STDY7770984 | 19/10/2017 | Busi | Intensive | Koome | ERS3016743 | pre_treatment |

**S2 Table.**
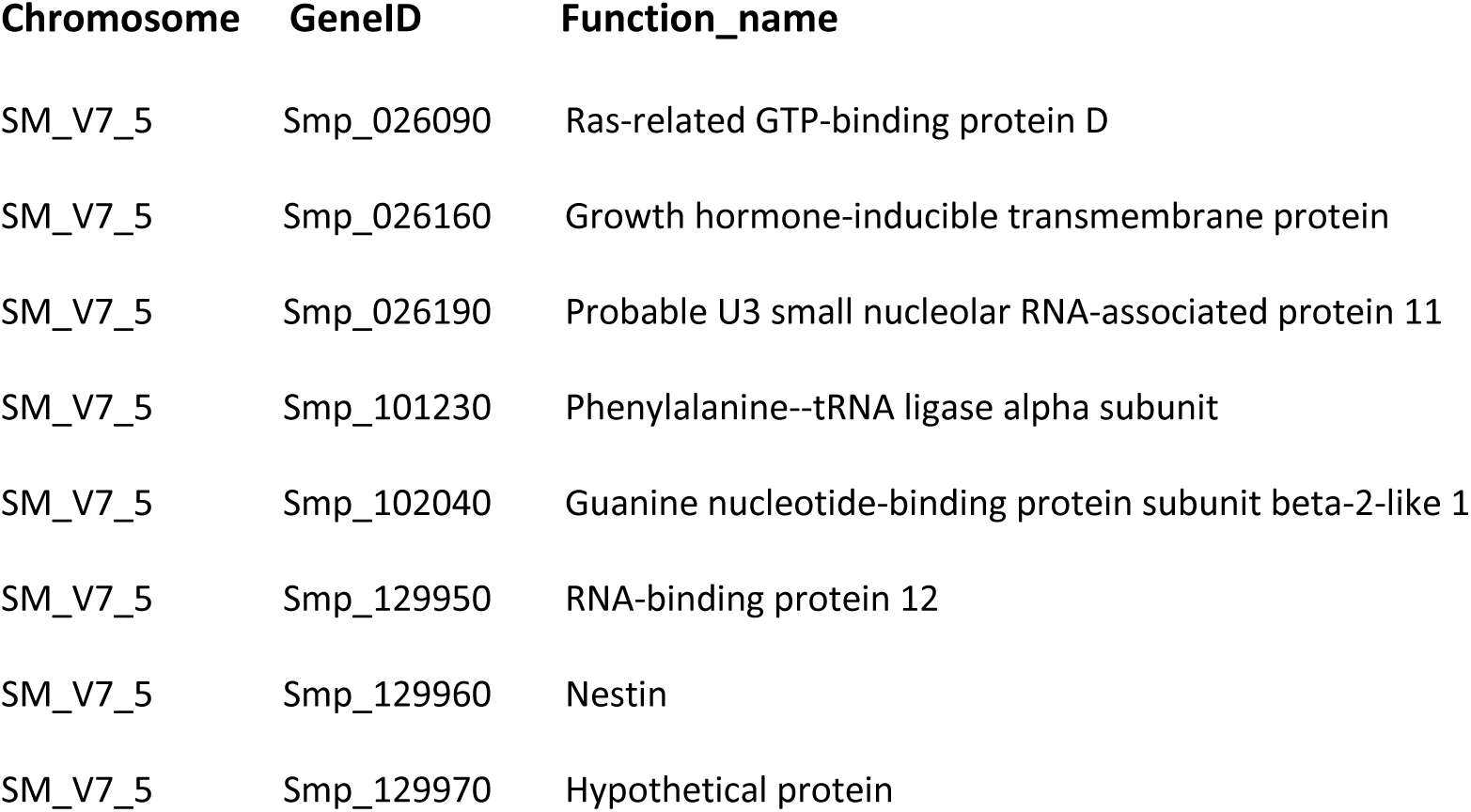
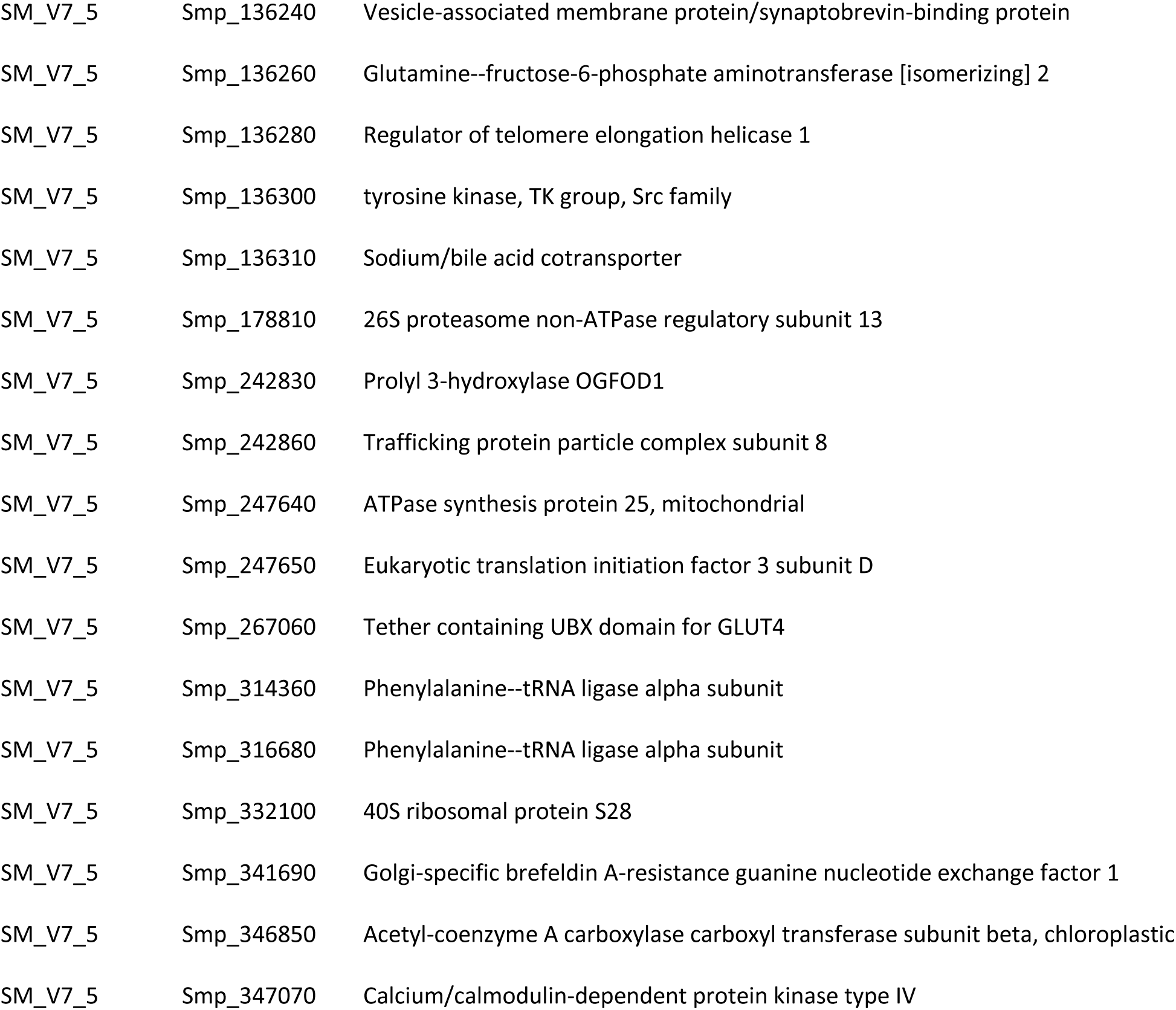
Protein coding genes present in the region of highest genetic differentiation on chromosome 5. (region X on Fig. 4B).

**S3 Table.**
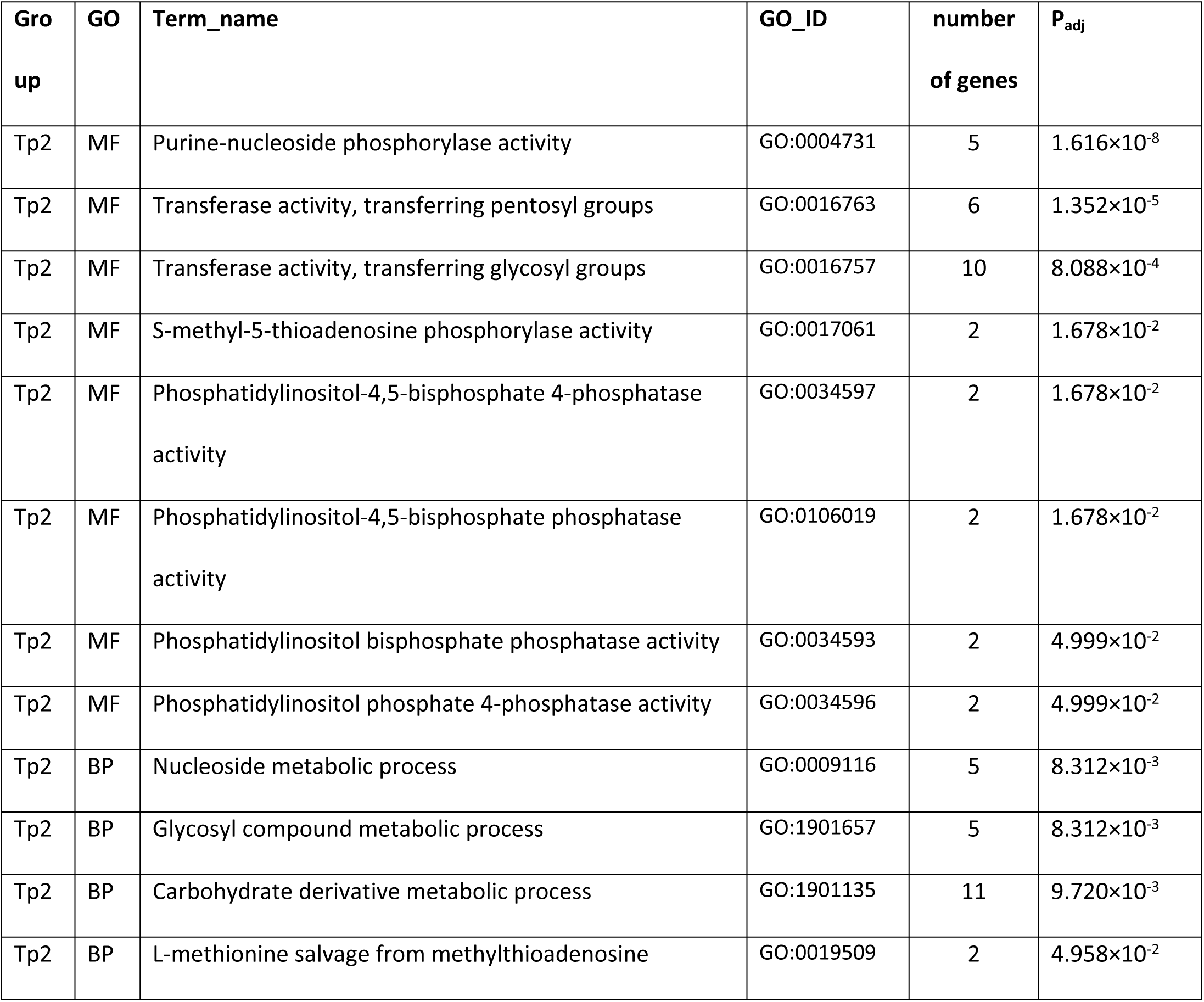
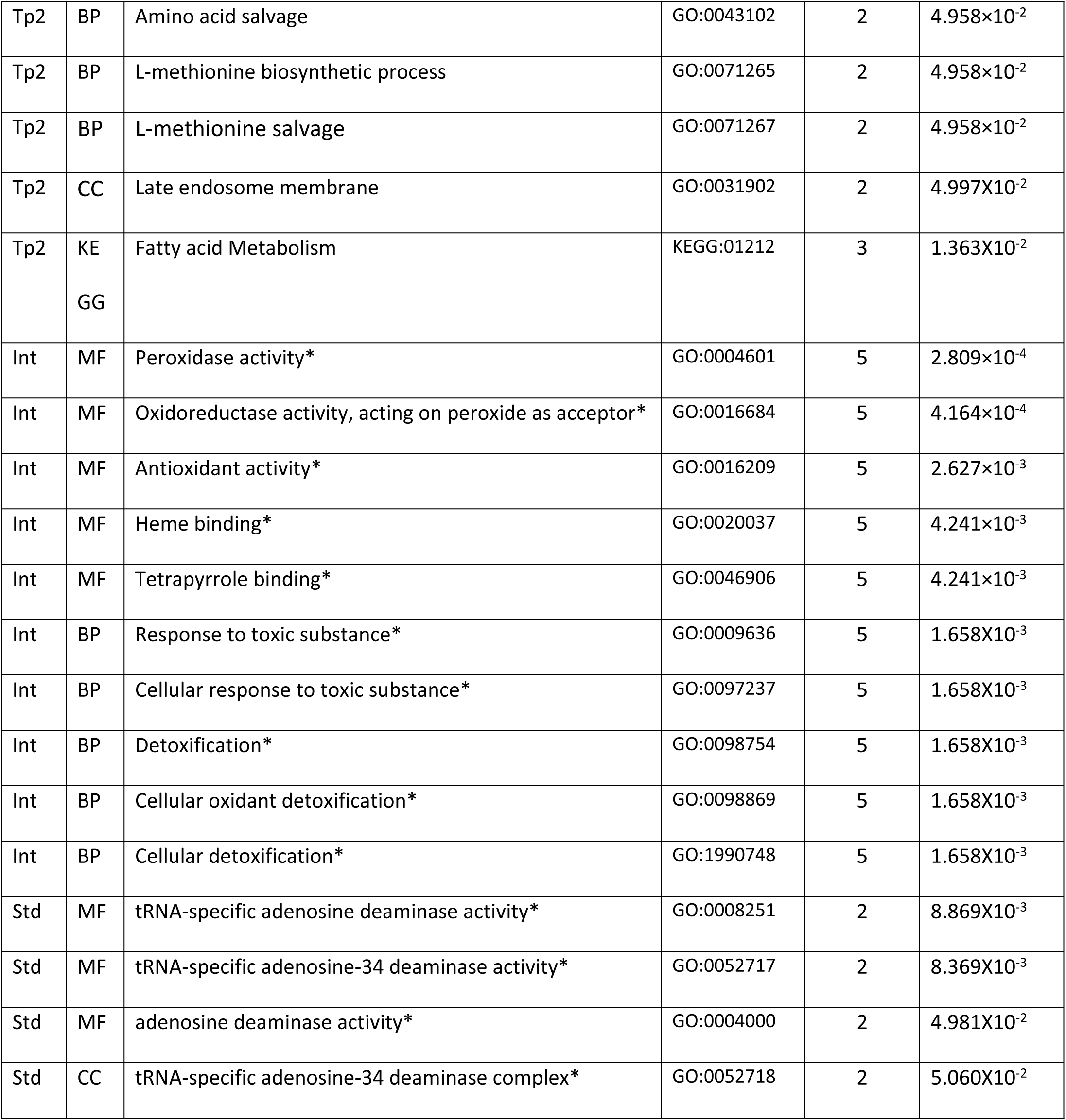
GO terms significantly over-represented among genes overlapping regions of different natural selection. Genes are from GO hierarchies for Molecular Function (MF), Biological Processes (BP), KEGG pathways and Cellular Component (CC) within the post-treatment (Tp2). Standard (Std) and Intensive (Int) groups with respective adjusted p-values (P_adj_). Asterisks mark terms that remain enriched in autosomal gene sets.

| Gro<br>up | GO | Term_name | GO_ID | number<br>of genes | $P_{adj}$ |
| --- | --- | --- | --- | --- | --- |
| Tp2 | MF | Purine-nucleoside phosphorylase activity | GO:0004731 | 5 | $1.616 \times 10^{-8}$ |
| Tp2 | MF | Transferase activity, transferring pentosyl groups | GO:0016763 | 6 | $1.352 \times 10^{-5}$ |
| Tp2 | MF | Transferase activity, transferring glycosyl groups | GO:0016757 | 10 | $8.088 \times 10^{-4}$ |
| Tp2 | MF | S-methyl-5-thioadenosine phosphorylase activity | GO:0017061 | 2 | $1.678 \times 10^{-2}$ |
| Tp2 | MF | Phosphatidylinositol-4,5-bisphosphate 4-phosphatase activity | GO:0034597 | 2 | $1.678 \times 10^{-2}$ |
| Tp2 | MF | Phosphatidylinositol-4,5-bisphosphate phosphatase activity | GO:0106019 | 2 | $1.678 \times 10^{-2}$ |
| Tp2 | MF | Phosphatidylinositol bisphosphate phosphatase activity | GO:0034593 | 2 | $4.999 \times 10^{-2}$ |
| Tp2 | MF | Phosphatidylinositol phosphate 4-phosphatase activity | GO:0034596 | 2 | $4.999 \times 10^{-2}$ |
| Tp2 | BP | Nucleoside metabolic process | GO:0009116 | 5 | $8.312 \times 10^{-3}$ |
| Tp2 | BP | Glycosyl compound metabolic process | GO:1901657 | 5 | $8.312 \times 10^{-3}$ |
| Tp2 | BP | Carbohydrate derivative metabolic process | GO:1901135 | 11 | $9.720 \times 10^{-3}$ |
| Tp2 | BP | L-methionine salvage from methylthioadenosine | GO:0019509 | 2 | $4.958 \times 10^{-2}$ |
| Tp2 | BP | Amino acid salvage | GO:0043102 | 2 | 4.958×10<sup>-2</sup> |
| Tp2 | BP | L-methionine biosynthetic process | GO:0071265 | 2 | 4.958×10<sup>-2</sup> |
| Tp2 | BP | L-methionine salvage | GO:0071267 | 2 | 4.958×10<sup>-2</sup> |
| Tp2 | CC | Late endosome membrane | GO:0031902 | 2 | 4.997X10<sup>-2</sup> |
| Tp2 | KE<br>GG | Fatty acid Metabolism | KEGG:01212 | 3 | 1.363X10<sup>-2</sup> |
| Int | MF | Peroxidase activity* | GO:0004601 | 5 | 2.809×10<sup>-4</sup> |
| Int | MF | Oxidoreductase activity, acting on peroxide as acceptor* | GO:0016684 | 5 | 4.164×10<sup>-4</sup> |
| Int | MF | Antioxidant activity* | GO:0016209 | 5 | 2.627×10<sup>-3</sup> |
| Int | MF | Heme binding* | GO:0020037 | 5 | 4.241×10<sup>-3</sup> |
| Int | MF | Tetrapyrrole binding* | GO:0046906 | 5 | 4.241×10<sup>-3</sup> |
| Int | BP | Response to toxic substance* | GO:0009636 | 5 | 1.658X10<sup>-3</sup> |
| Int | BP | Cellular response to toxic substance* | GO:0097237 | 5 | 1.658X10<sup>-3</sup> |
| Int | BP | Detoxification* | GO:0098754 | 5 | 1.658X10<sup>-3</sup> |
| Int | BP | Cellular oxidant detoxification* | GO:0098869 | 5 | 1.658X10<sup>-3</sup> |
| Int | BP | Cellular detoxification* | GO:1990748 | 5 | 1.658X10<sup>-3</sup> |
| Std | MF | tRNA-specific adenosine deaminase activity* | GO:0008251 | 2 | 8.869X10<sup>-3</sup> |
| Std | MF | tRNA-specific adenosine-34 deaminase activity* | GO:0052717 | 2 | 8.369X10<sup>-3</sup> |
| Std | MF | adenosine deaminase activity* | GO:0004000 | 2 | 4.981X10<sup>-2</sup> |
| Std | CC | tRNA-specific adenosine-34 deaminase complex* | GO:0052718 | 2 | 5.060X10<sup>-2</sup> |

## Notes

### Competing Interest Statement

The authors have declared no competing interest.

